# Imaging the Neural Substrate of Trigeminal Neuralgia Pain Using Deep Learning

**DOI:** 10.1101/2022.11.02.514527

**Authors:** Yun Liang, Qing Zhao, Zhenhong Hu, Ke Bo, Sreenivasan Meyyappan, John K. Neubert, Mingzhou Ding

## Abstract

Trigeminal neuralgia (TN) is a severe and disabling facial pain condition and is characterized by intermittent, severe, electric shock-like pain in one (or more) trigeminal subdivisions. This pain can be triggered by an innocuous stimulus or can be spontaneous. Presently available therapies for TN include both surgical and pharmacological management; however, the lack of a known etiology for TN contributes to the unpredictable response to treatment and the variability in long-term clinical outcomes. Given this, a range of peripheral and central mechanisms underlying TN pain remain to be understood. We acquired functional magnetic resonance imaging (fMRI) data from TN patients who (1) rested comfortably in the scanner during a resting state session and (2) rated their pain levels in real time using a calibrated tracking ball-controlled scale in a pain tracking session. Following data acquisition, the data was analyzed using the conventional correlation analysis and two artificial intelligence (AI)-inspired deep learning methods: convolutional neural network (CNN) and graph convolutional neural network (GCNN). Each of the three methods yielded a set of brain regions related to the generation and perception of pain in TN. There were six regions that were identified by all three methods, including the superior temporal cortex, the insula, the fusiform, the precentral gyrus, the superior frontal gyrus, and the supramarginal gyrus. Additionally, 17 regions, including dorsal anterior cingulate cortex(dACC) and the thalamus, were identified by at least two of the three methods. Collectively, these 23 regions represent signature centers of TN pain and provide target areas for future studies relating to central mechanisms of TN.

## Introduction

Trigeminal neuralgia (TN) is a severe and disabling facial pain condition and is considered one of the most, if not the most, painful conditions a person can endure. TN is characterized by severe, intermittent, paroxysmal, electric shock-like pain attacks in one (or more) trigeminal subdivision and this pain can be triggered by an innocuous stimulus or can be spontaneous. The 3rd edition of the International Classification of Headache Disorders (ICHD-3) classifies TN into idiopathic, classical, or secondary trigeminal neuralgia diagnoses.^1^ Idiopathic TN, by definition, has no known etiology, while classical TN is a diagnosis given when there is verification (e.g., visualization during surgery and/or via neuroimaging) of blood vessel contact against the root of the trigeminal nerve on the ipsilateral side of the pain complaint. Secondary TN is caused by an identifiable pathological condition, including multiple sclerosis or tumor impingement on the trigeminal root trunk, most commonly at the cerebellopontine angle. For classical and idiopathic TN, recent diagnostic subdivisions also include either (1) a purely paroxysmal quality or (2) having intermittent paroxysmal bouts with a more continuous background of burning and/or aching pain.^1,2^ Despite the detailed criteria for diagnosing the various TN disorders, little is known regarding etiology, in particular what central mechanisms may contribute to this pain condition.^3^ For example, while classical TN is proposed to have a neurovascular insult as the etiology, many cases of TN do not show this impingement; conversely, many patients have impingement, but no pain. Thus, the distinction between classical and idiopathic is more semantics rather than explanatory in regard to the distinct pain symptoms associated with TN.

First line treatments of TN involve medication trials with anti-convulsant medications such as carbamazepine and oxcarbazepine. In fact, responding to carbamazepine is pathognomonic for TN. Anticonvulsant medications often have intolerable side effects and become progressively less effective with time.^4^ Following failure of medication trials, patients usually escalate treatment to invasive surgical procedures. The most common surgical procedure is called microvascular decompression (MVD), which involves moving an impinging blood vessel (e.g., superior cerebellar artery) off the trigeminal nerve and placing a Teflon pad to maintain this neurovascular separation. While surgical procedures provide lasting relief for some patients, for others, the relief can be short-lived, and pain returns after a few months to a few years. According to available data, approximately 4% of patients per year experience recurrence of TN pain after MVD,^5^ and over a period of 10-20 years, the recurrence rate may exceed 10%.^6^ While direct visualization during surgery provides verification of the classical TN diagnosis, there are TN patients who do not respond to surgery. Additionally, there is a cohort of TN patients who do not have this vascular insult of the trigeminal nerve, thus indicating a range of peripheral and central mechanisms underlying TN pain remain to be understood.

Functional magnetic resonance imaging (fMRI) remains the main neuroimaging technique for investigating the central mechanisms of pain. In particular, resting-state fMRI (rs-fMRI), which measures blood-oxygen-level-dependent (BOLD) signals while the patient is not engaged in any systematic thought or activity, has been applied extensively in TN.^7,8^ Compared to age-and sex-matched healthy control subjects, Yuan et al.^7^ found that TN patients exhibited significantly increased regional homogeneity (ReHo) and signal amplitude in several brain regions, including the posterior lobe of the cerebellum, anterior cingulate cortex, middle temporal gyrus, temporal lobe, putamen, occipital lobe, limbic lobe, precuneus, and the medial and superior frontal gyrus, and decreased ReHo in insula. They suggested that these abnormal activities played a role in the development and maintenance of chronic TN pain. Wang et al.^9^ found increased ReHo in the inferior temporal gyrus, thalamus, inferior parietal lobule, precentral and postcentral gyri and decreased ReHo in the amygdala, parahippocampal, and cerebellum in TN patients. Meanwhile, Yan et al.^10^ detected that TN patients had decreased ReHo in the left middle temporal gyrus, superior parietal lobule, and precentral gyrus and increased ReHo in the thalamus in the resting state studies, which also provided compelling evidence for abnormal resting-state brain activity in TN. Moving beyond regional analysis, at the network level, Zhu et al.^11^ applied functional connectivity methods to show that TN patients exhibited significantly higher degree centrality values in the right lingual gyrus, right postcentral gyrus, left paracentral lobule, and bilateral inferior cerebellum. They proposed that these changes reflect the adaptation of the cerebral cortex to frequent pain attacks over a long period of time. In addition to resting-state fMRI, Moisset et al.^12^ evoked pain in TN patients with stimulation of the cutaneous trigger zone and found increased activity in postcentral and precentral cortex, contralateral supplementary motor area, thalamus, anterior and posterior insula, prefrontal cortex, putamen, ipsilateral midcingulate cortex, hippocampus/parahippocampal area and cerebellum. **Table 1** summarizes these neuroimaging studies, highlighting the brain regions identified as having abnormal neural activities in TN subjects. Note that, with the exception of the cerebellum, there is a relative lack of consistency for identified brain structures across the studies, suggesting additional studies are needed for understanding and identifying brain structures that are functionally important in the generation and maintenance of TN pain.

**Table 1:** Regions showing abnormal activities in TN per published literature.

| Brain Areas | Yuan et al.<br>(2018) | Wang et al.<br>(2015) | Zhu et al.<br>(2020) | Yan et al.<br>(2019) | Moisset et al.<br>(2012) |
| --- | --- | --- | --- | --- | --- |
| Cerebellum | ✓ | ✓ | ✓ |  | ✓ |
| Cingulate cortex | ✓ |  |  |  | ✓ |
| Putamen | ✓ |  |  |  | ✓ |
| Middle temporal gyrus | ✓ |  |  | ✓ |  |
| Precuneus | ✓ |  |  |  |  |
| Medial frontal gyrus | ✓ |  |  |  |  |
| Superior frontal gyrus | ✓ |  |  |  |  |
| Thalamus |  | ✓ |  | ✓ | ✓ |
| Parietal lobule |  | ✓ |  | ✓ |  |
| Postcentral gyrus |  | ✓ | ✓ |  | ✓ |
| Amygdala |  | ✓ |  |  |  |
| Parahippocampal |  | ✓ |  |  | ✓ |
| Inferior temporal gyrus |  | ✓ |  |  | ✓ |
| Lingual gyrus |  |  | ✓ |  |  |
| Paracentral lobule |  |  | ✓ |  |  |
| Fusiform gyrus |  |  |  |  |  |
| Middle occipital gyrus |  |  |  |  |  |
| Precentral gyrus |  |  |  | ✓ | ✓ |
| Secondary somatosensory cortex |  |  |  |  | ✓ |
| Supplementary motor area |  |  |  |  | ✓ |
| Superior temporal gyrus |  |  |  |  | ✓ |
| Insula |  |  |  |  | ✓ |

Chronic pain can cause changes in many structures in the brain.^13^ While the neural activity in a brain area may appear to be altered in rs-fMRI studies, it may not imply that the area is directly involved in pain generation or perception. Additionally, stimulus-evoked pain may differ fundamentally from spontaneous or continuous pain, as experienced with TN.^14^ To address this issue, a neuroimaging technique that directly addresses the neural substrate of the naturally occurring pain is percept-related fMRI.^15^ In this technique the patients indicate their moment-to-moment pain levels while their brain activities are being recorded.^16^ Correlating brain activity and pain level fluctuations one obtains information on brain structures that directly underlie pain generation and perception. Kwan et al.^17^ applied this percept-related fMRI technology to patients with irritable bowel syndrome (IBS) and revealed abnormal urge- and pain-related forebrain activity during rectal distension. Baliki et al.^18^ applied the method to patients with chronic back pain (CBP) and found that the insular region was active when pain level increases. To date, no studies have applied the percept-related fMRI approach to investigate TN pain.

We recorded fMRI data while TN patients (1) rated their spontaneous pain levels in the pain tracking session and (2) rested in the resting state session. The data were first analyzed using the conventional correlation method, which assessed whether fMRI signals from different brain regions were correlated with the fluctuating pain ratings, with brain regions showing strong correlations considered the “signature” centers of TN pain. In addition, we also applied deep neural networks (DNNs), a branch of deep learning, to further interrogate the data.^19^ DNNs, including convolution neural networks (CNNs) and graph convolution neural networks (GCNNs), differ from traditional machine learning techniques such as support vector machine (SVM) and logistic regression (LR) in that the hidden layers in these models are capable of encoding and utilizing more complex features of the data to provide more accurate predictions of experimental conditions and a deeper understanding of the data.^20^ While SVM and LR have been applied to chronic pain conditions, e.g. to differentiate healthy control subjects from patients with fibromyalgia and CBP,^21,22^ DNNs have only begun to see applications in chronic pain studies.^23^ The present study is the first to train DNN models to predict the fluctuating pain ratings from fMRI data in TN and to derive the signature centers of TN pain from these models. Our overall goal is to seek converging evidence on the neural substrate of TN pain by combining different methods and to uncover novel insights not possible with the conventional methods.

## Materials and Methods

The study was approved by the WCG Institutional Review Board (IRB). Patients were recruited through the clinical care population within the University of Florida Health System and from referrals provided by the Facial Pain Research Foundation (FPRF). Screening was done either in person or via a phone call. Eligible subjects were consented and completed a study packet that included a Health History Questionnaire, Oregon Health Science University (OHSU) Trigeminal Neuralgia – Diagnostic Questionnaire, Beck Depression Inventory-II (BDI-2), Pain Anxiety Screening Scale (PASS), and Pain Catastrophizing Scale (PCS). The study coordinator read a prepared standard script explaining the study procedures. A focused medical history, a trigeminal cranial nerve exam, and a physical exam was completed by a trained clinical fellow and the PI (JN). Vital signs (blood pressure, temperature, and pulse) were also recorded prior to scanning.

### Pain ratings

Subjects indicated on a 100mm visual analog scale (VAS) anchored on the left with “no pain sensation” and on the right with “most intense pain sensation imaginable” their daily experienced pain (past month). This rating is henceforth referred to as “usual pain”. Subjects also used the VAS to rate their “current pain” just prior to entering the MRI scanner. During the scanning procedure, subjects completed continuous pain tracking (see Experimental Paradigm below, **Figure 1 (B)**) where they could visualize a computer screen in the scanner via mirrors and rate their pain levels in real time using a tracking ball.

**Figure 1.**
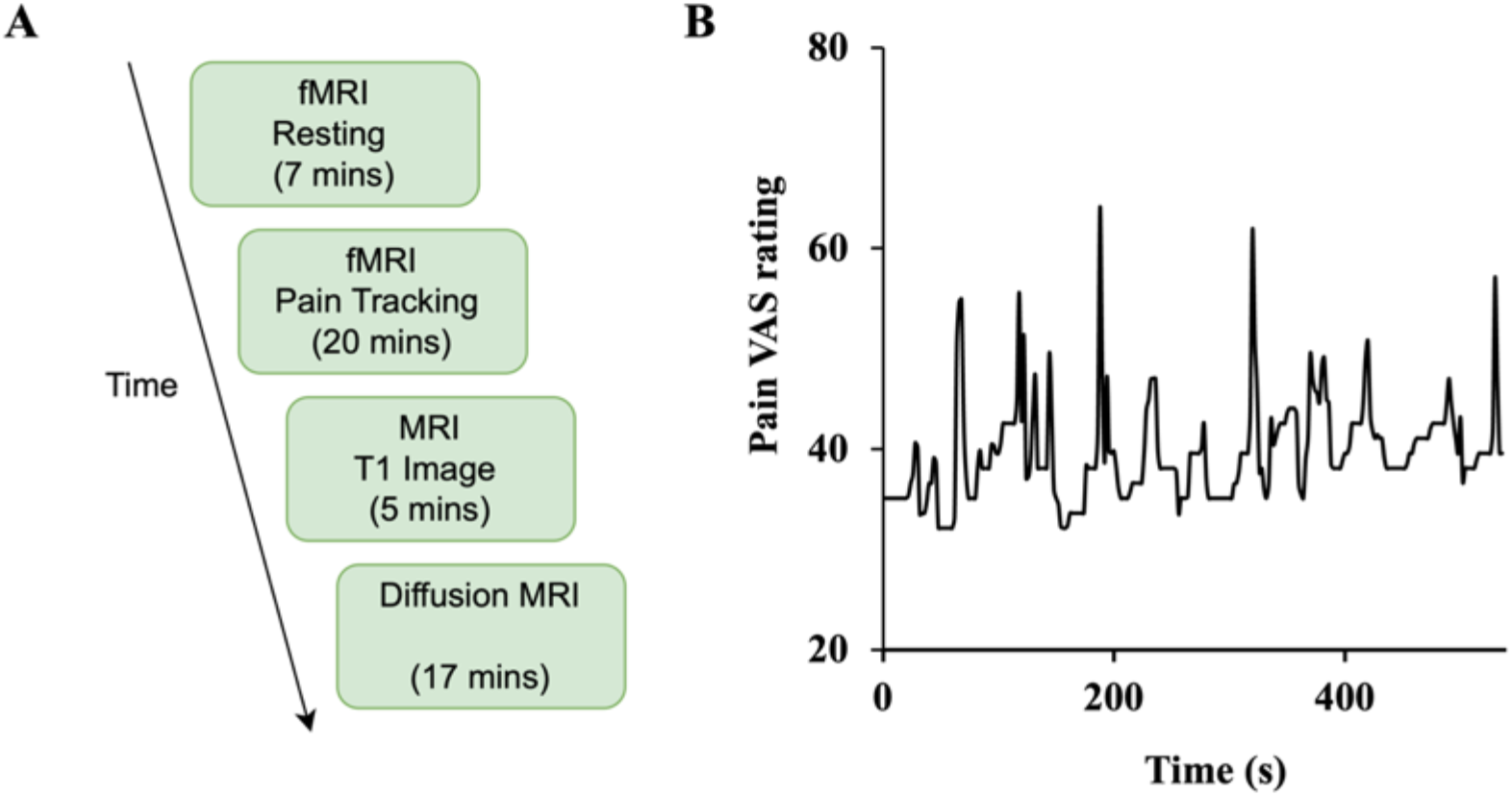
Study design. **(A)** The experiment was divided into four sessions: resting-state (functional), pain tracking (functional), T1 imaging (structural), and diffusion MRI (structural). **(B)** Sample time course of spontaneous ratings of pain from a representative patient.

### Participants

#### Inclusion Criteria

- Male and female subjects
- Age 18-75 years old. This represents the age range of many chronic pain patients, including patients with TN.
- American Society of Anesthesiologists (ASA) status 1, 2, or 3, deemed in good general health.
- Only subjects with reported average usual pain of moderate to severe (VAS of 30-100 mm) at the time of screening were included.
- Subjects were diagnosed with trigeminal neuralgia, per the International Headache Society (IHS) Disorders criteria,^1^ as having:

A. Recurrent paroxysms of unilateral facial pain in the distribution(s) of one or more divisions of the trigeminal nerve, with no radiation beyond, and fulfilling criteria B and C.
B. Pain has all of the following characteristics:
  1. lasting from a fraction of a second to two minutes
  2. severe intensity
  3. electric shock-like, shooting, stabbing or sharp in quality
C. Precipitated by innocuous stimuli within the affected trigeminal distribution.
D. Not better accounted for by another ICHD-3 diagnosis.
- Subjects included having symptoms as purely paroxysmal pain or having intermittent paroxysmal bouts with a more continuous background of burning and/or aching pain.

Note that during the screening process, individuals reported being diagnosed with TN by their physician would have this diagnosis verified by the study staff. Additionally, subjects completed the OHSU TN – Diagnostic Questionnaire as additional verification of TN diagnosis.

#### Exclusion Criteria

- Patients diagnosed with trigeminal neuralgia attributed to space-occupying lesion (ICHD-3 code: 13.1.1.2.2) or other cause (e.g., multiple sclerosis, ICHD-3 code: 13.1.1.2.3), painful trigeminal neuropathy (ICHD-3 code: 13.1.2), trigeminal post-herpetic neuralgia (ICHD-3 code: 13.1.2.2), trigeminal neuropathic pain (ICHD-3 code: 13.1.2.4), and idiopathic painful trigeminal neuropathy (ICHD-3 code: 13.1.2.5).
- ASA status 4-5 and Emergency operation.
- Presence of chronic disease (*e*.*g*. cardiovascular disease, liver disease, kidney disease, diabetes, etc.), other than trigeminal neuralgia.
- Pregnant females.
- No exclusions were made based on race, gender, or religion.

In total, 55 patients gave written informed consent and participated in the study (69% female, mean age ± standard deviation (SD)=53.3±14.9). Sixteen patients were rejected due to a combination of the following reasons: (1) not meeting diagnosis criteria (*n=1*), (2) not completing the whole experiment (*n=11*), (3) technical difficulties during fMRI recording (*n=2*), and (4) excessive movements inside the scanner (*n=2*). The data from the remaining 39 patients were analyzed and reported here. Of the 39 patients, diagnostic concordance between the PI and the OHSU Trigeminal Diagnostic Questionnaire was 38/39 (97.5%). For the 1/39 patient, the OSHU TN diagnosis was nervus intermedius neuralgia, which is characterized by an intermittent stabbing deep pain in the ear with associated tinnitus. However, during the examination, it was determined that this one subject met the inclusion criteria for idiopathic TN, having intermittent paroxysmal bouts with a more continuous background of burning and/or aching pain. The vital signs taken just prior to scanning were within normal limits for all subjects (data not shown) and no adverse reactions or events were reported during any of the procedures.

### Experimental paradigm

Subjects underwent functional, structural, and diffusion magnetic resonance imaging. As shown in **Figure 1 (A)**, there are two types of functional scans: resting state and pain tracking. During the resting state scan (seven minutes), the patients were instructed to fixate on the cross at the center of the monitor screen, stay still, and not think any systematic thought. During the pain tracking scan (10-20 minutes), fMRI data were acquired while the patients rated their momentary pain levels using a tracking ball. The tracking ball controlled the movement of a cursor along a straight line with 0 and 100 indicated at the two ends of the line on the monitor. An example of a pain tracking time course from one patient is shown in **Figure 1 (B)**.

For the first 10 subjects, the pain tracking session was divided into two parts. For the first 10 minutes the subjects tracked their pain level fluctuations as described above. The second 10 minutes was a motion tracking session in which a marker moved on the monitor between 0 and 100 according to the subject-indicated pain level fluctuations from the previous pain tracking session. The subject was asked to move the tracking ball to track the movement of the marker. We had to discontinue the motion tracking session after the first 10 subjects because it was becoming apparent that 10 minutes of actual pain tracking was not enough to produce sufficient data for the intended analyses. For the remaining patients the pain tracking session lasted the entire 20 minutes. It is worth noting that increasing the length of the overall experiment was not an option because of the burden it would place on the patient.

### Data acquisition and preprocessing

Functional MRI images were collected on a 3T Philips Achieva scanner (Philips Medical Systems, the Netherlands) equipped with a 32-channel head coil. The echo-planar imaging (EPI) sequence parameters were as follows: repetition time (TR), 1.98 sec; echo time, 30 msec; flip angle, 80; field of view, 224 mm; slice number, 36; voxel size, 3.5×3.5×3.5 mm; matrix size, 64×64. In addition to the functional scans, a high-resolution anatomical T1-weighted MRI image was also acquired for each subject using the following parameters: field of view (FOV), 240×240 mm; time of repetition (TR), 8.0566 ms; time of echo (TE), 3.686 ms; resolution, 1×1 mm; flip angle, 80 degrees.

Statistical parametric mapping (SPM) was used to preprocess the functional MRI data.^24^ The preprocessing steps include slice timing correction, realignment, co-register, spatial normalization, and spatial smoothing. Data segments with large head movements were removed from five subjects. For the fMRI data that were subjected to further analysis, head motions were regressed out, and a bandpass filter [0.01, 0.1 Hz] was applied to reduce low-frequency and high-frequency noise.

### Correlation analysis

This analysis has been applied in prior percept-related fMRI studies of chronic pain. In this analysis, the time course of pain ratings (see, for example, **Figure 1 (B))** was first convolved with the hemodynamic response function (HRF), and then correlated with the BOLD time course from every voxel in the brain.^25^ The correlation values were Fisher-transformed to achieve approximate normal distribution and subjected to the population level analysis. A threshold was set at *P<0*.*05* uncorrected. To reduce the influence of possible spurious correlations, only voxels that are part of clusters of at least 10 voxels meeting this statistical threshold were considered in the brain map. Both positively correlated voxels and negatively correlated voxels were considered. Regions showing strong correlations were considered as potential signature centers of TN pain. This pipeline of the correlation analysis is illustrated in **Figure 2**.

**Figure 2.**
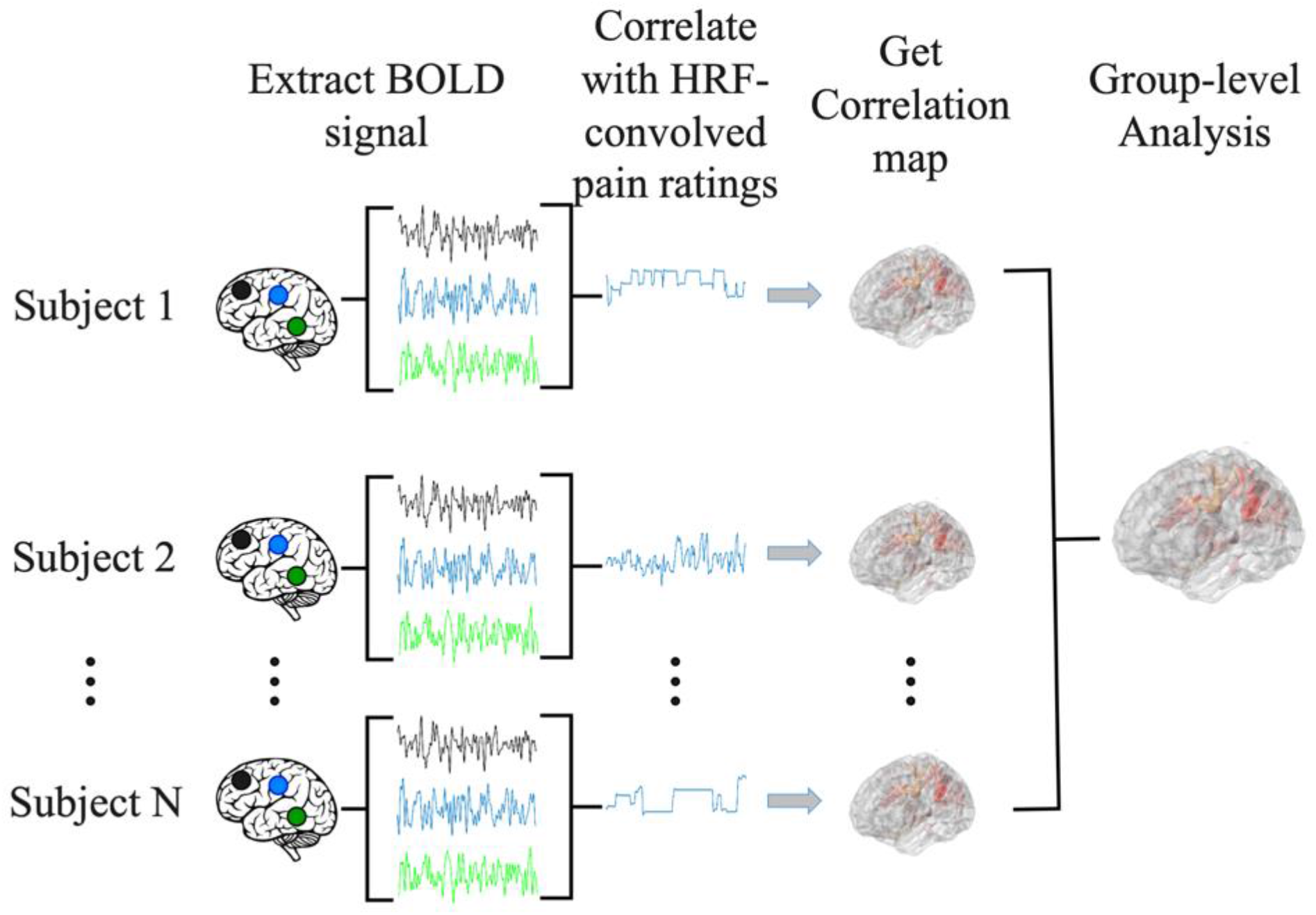
Pipeline for correlation analysis. BOLD signal from each voxel was correlated with the hemodynamic response function (HRF)-convolved pain rating time course to generate a correlation map for each subject. A group level analysis was then performed to generate the population level correlation map.

### CNN analysis

Increasingly, artificial intelligence (AI)-inspired techniques such as DNNs are being applied to analyze fMRI data, providing insights not possible with other techniques.^26^ We implemented a CNN model, which is a type of DNN, to predict pain ratings from fMRI data. Given that CNNs are more adapted to predict discrete labels rather than continuous values, we divided continuous pain ratings into two categories: low pain (pain ratings<=15) and high pain (pain ratings>15). Here the threshold of 15 was chosen so that the number of data points in the high and low pain categories were approximately equal across the entire patient population. Numerically, low pain has a value of zero whereas high pain has a value of one. The CNN, as shown **Figure 3 (A)**, consisted of ResNeXt101 with four cardinalized res-blocks with 32 independent paths within each block.^27^ The convolutional filters were modified and made three-dimensional so that they can be applied to the three-dimensional fMRI data. The final output layer contained two units corresponding to low pain and high pain respectively. The 39 patients were divided into eight groups of four-five patients each. Seven groups were chosen as the training dataset and the remaining group was chosen as the testing dataset. This process was repeated eight times (eight-fold cross validation). The reported decoding accuracy and the receiver operating characteristic (ROC) curve were the averages from the eight repetitions. For model training, the weighted cross-entropy was used as the loss function and the stochastic gradient descent (SGD) as the optimizer, with a momentum of 0.9 and a weight decay of 5.0e-4. The initial learning rate was set as 1.0e-3. The number of training epochs was 50 with a batch size of two.

**Figure 3.**
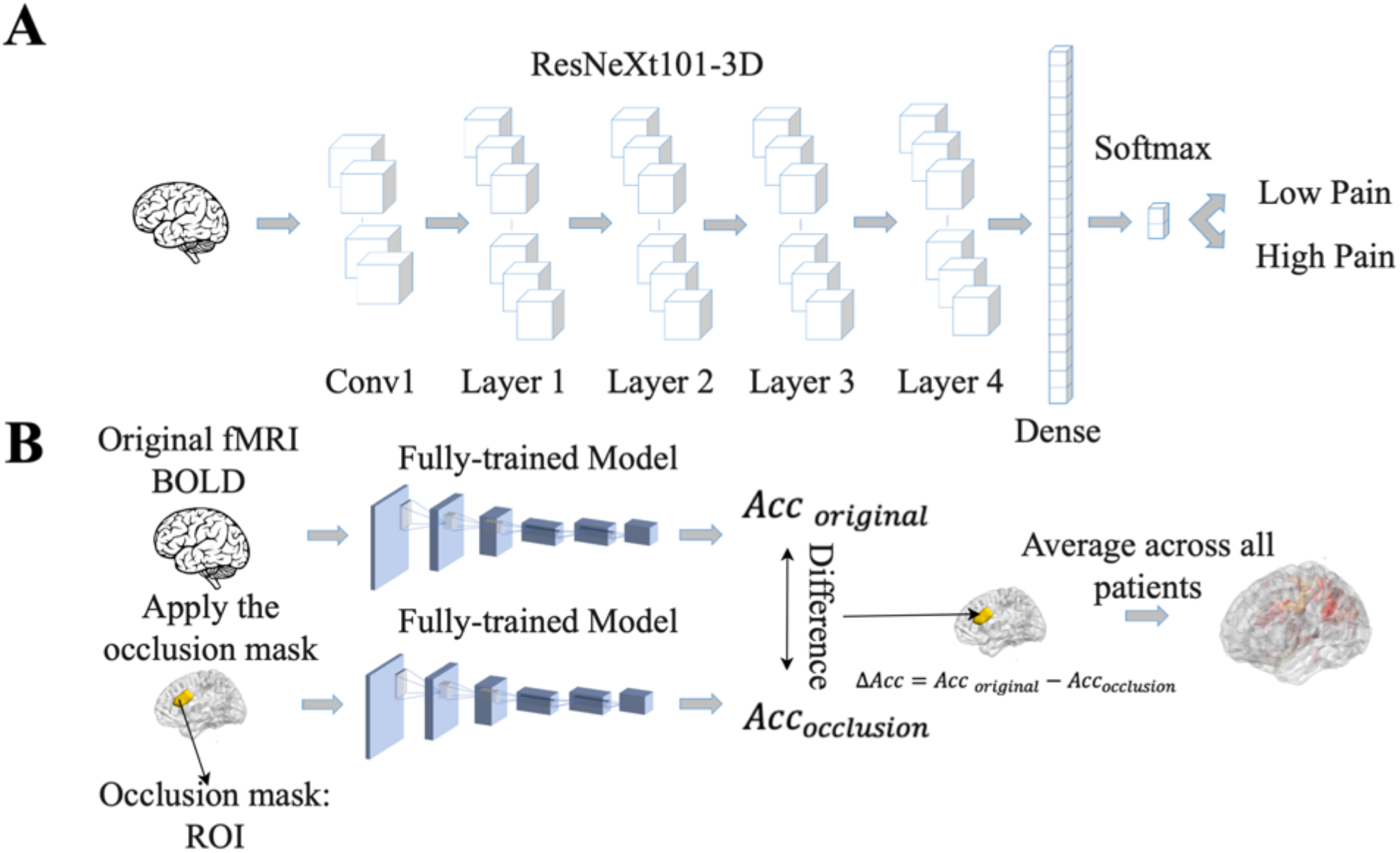
Pipeline for Convolutional Neural Network (CNN) analysis. **(A)** CNN model was trained to predict pain rating fluctuations from fMRI data. **(B)** An occlusion approach was carried out to evaluate the contribution of different brain regions to CNN model prediction performance. Acc: accuracy.

Once the CNN model was shown to have the ability to predict pain ratings from fMRI data, we proceeded to identify the essential brain regions that contribute to the prediction performance. The occlusion method was used for this purpose. See **Figure 3 (B)**. For each region of interest (ROI) in the Lausanne atlas,^28^ the BOLD values in all of its voxels were replaced by zero and fed into the CNN model, and the decoding accuracy was recorded. The amount of decoding accuracy decrease compared to the full data decoding accuracy was taken as a measure of the importance of the ROI in model prediction. The more the prediction accuracy decreases from occluding a brain region, the more that brain region contributes to the prediction performance, and the more weight it gets in the resulting heatmap. The reported heatmaps were the average from the eight models described earlier.

### GCNN analysis

It is increasingly recognized that pain processing involves multiple brain areas and their interactions and is a network phenomenon.^29^ The correlation analysis and the CNN analysis described above do not take into account functional relationships between different brain regions during pain processing. To address this problem, we implemented a GCNN approach to predict pain ratings from fMRI data. The GCNN, shown in **Figure 4 (A)**, consisted of two GIN layers,^30^ and one fully connected layer containing two output units for predicting low pain vs high pain. The Lausanne atlas was used to divide the brain into 129 ROIs.^28^ BOLD signals within each ROI were averaged and ROI-to-ROI dynamic interactions were assessed by cross-correlation in moving windows of 30s in duration. After the cross-correlation matrices were computed, a binarization was carried out by applying a threshold, where the values greater than the threshold were set to one and smaller than the threshold were set to zero. The binarized connectivity matrix along with the average BOLD signals from each ROI were taken as input features for the GCNN to predict the pain ratings in the middle of the same 30s moving window. The patients were again divided into eight groups of four-five each (same eight-fold validation as CNN). We applied the weighted cross-entropy as our loss function and SGD as our optimizer with a momentum of 0.9 and a weight decay of 5.0e-4 during the model’s training phase. The initial learning rate was set as 1.0e-3. The reported decoding accuracy and the ROC curve were averages of the eight-fold results.

**Figure 4.**
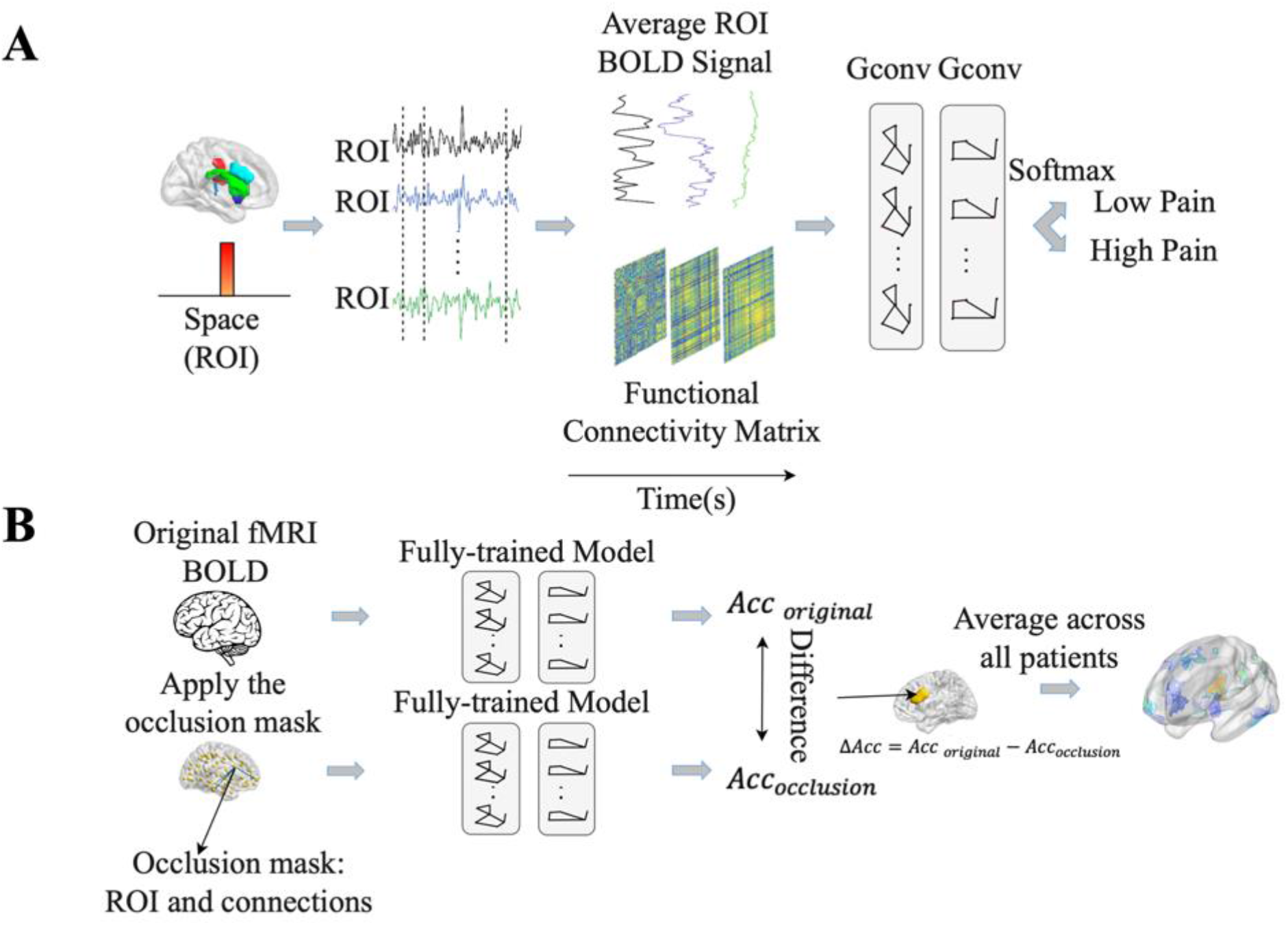
Pipeline for Graph Convolutional Neural Network (GCNN) analysis. **(A)** GCNN model was trained to predict pain rating fluctuations from fMRI data. **(B)** An occlusion approach was carried out to evaluate the contribution of different brain regions and their associated functional connections to GCNN model prediction performance. Acc: accuracy.

Once the models were shown to have the ability to predict pain ratings from fMRI data, we proceeded to apply the occlusion method to identify essential network nodes through which the functional interactions among different brain areas play an essential role in pain prediction. Each ROI along with the connections through the ROI were replaced by zero and the prediction performance decline was calculated. The degree of decline is taken to indicate the importance of the ROI in mediating pain related network processing. A heatmap was derived based on this principle.

### Data availability

The datasets for this article are not publicly available because other studies have been conducted with the same data and they are not published yet. Requests to access the datasets should be directed to the PI at.

## Results

We recorded fMRI data while *n=39* TN patients (1) tracked their pain level fluctuations in the pain tracking session and (2) rested without thinking any systematic thought in the resting state session. Both conventional correlation analysis and AI-inspired analyses (CNN and GCNN) were applied to reveal the signature centers in the brain underlying the generation and perception of TN pain.

### Correlation analysis

The data from the pain tracking session was analyzed here. The BOLD time course from each voxel was correlated with the HRF-convolved pain rating time course. A voxel was considered potentially part of the neural substrate underlying TN pain if this correlation is strong regardless of the sign of correlation. For the subject in **Figure 1 (B)**, the HRF-convolved pain rating time course had a strong positive correlation with the BOLD signal from a voxel located in the precentral gyrus (**Figure 5 (A)**, *R=0*.*55*), a low correlation with the BOLD signal from a voxel located in the superior frontal cortex (**Figure 5 (B)**, *R=-0*.*03*), and a strong negative correlation with the BOLD signal from another voxel located in the superior frontal cortex (**Figure 5 (C)**, *R=-0*.*61*). Across all patients, using uncorrected *P<* 0.05 and a minimum cluster size of 10 voxels as the criteria, a set of brain regions showing both strong positive and negative correlations with pain were identified (**Figure 5 (D)** and **Figure 5 (E))**, including the precentral gyrus, the lingual, the thalamus, the superior frontal, and the superior temporal. See **Tables 2** and **3**. Some regions may have more than one cluster of correlated voxels. In addition, given that the ROIs are relatively large, positive correlation clusters and negative correlation clusters may appear in the same ROI (e.g., superior frontal). Combining **Tables 2** and **3**, there are 21 distinct ROIs that showed strong correlations with spontaneous pain fluctuations.

**Figure 5.**
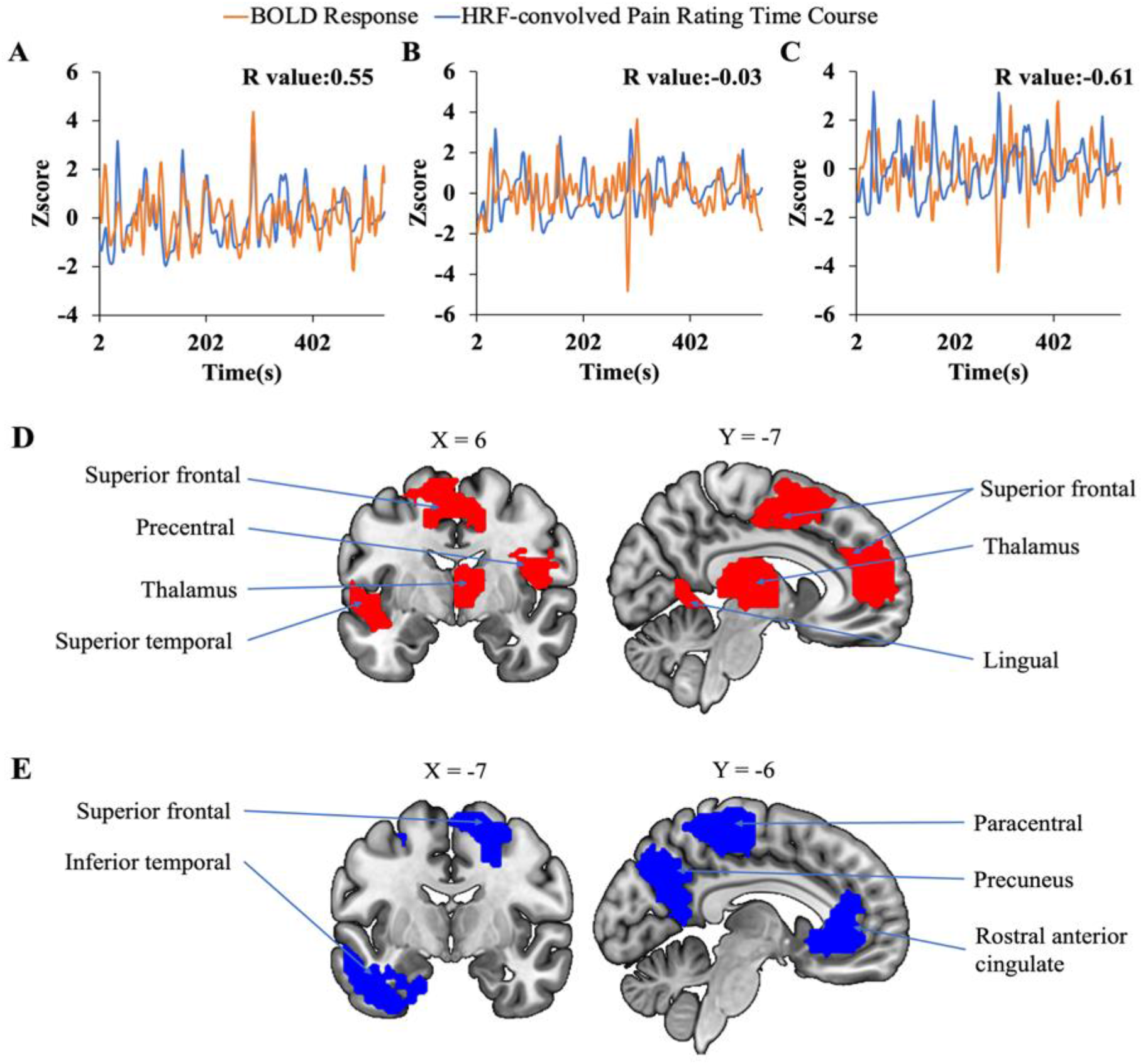
Correlation analysis. **(A)-(C)** Correlation between voxel-level fMRI time series and HRF convolved pain ratings. The HRF-convolved pain ratings had a strong correlation (*R = 0*.*55*) with the BOLD signal from one voxel in the precentral gyrus (A), a low correlation (*R = −0*.*03*) with the BOLD signal from a voxel located in the superior frontal gyrus (B), and a strong negative correlation (*R = −0*.*61*) with the BOLD signal from one voxel located in the superior frontal cortex (C). **(D)-(E)** Regions showing strong positive (D) and negative (E) correlation with pain ratings.

**Table 2:** Regions showing positive BOLD-pain correlations.

| Region | Coordinates contrast |  |  | Volume(voxel) | Volume(mm<sup>3</sup>) | peak T score |
| --- | --- | --- | --- | --- | --- | --- |
| Superior frontal | 3 | 44 | 31 | 203 | 5481 | 3.74 |
|  | -27 | -7 | 61 | 60 | 1620 | 2.34 |
|  | 15 | 65 | 7 | 14 | 378 | 2.1 |
|  | 6 | -4 | 49 | 110 | 2970 | 2.08 |
| Lateral orbitofrontal | 30 | 20 | -11 | 97 | 2619 | 3.14 |
|  | -33 | 20 | -11 | 18 | 486 | 2.22 |
| Precentral | 54 | 14 | 25 | 40 | 1080 | 2.76 |
|  | -33 | -25 | 49 | 28 | 756 | 2.41 |
| Thalamus | 12 | -10 | 16 | 41 | 1107 | 2.64 |
|  | 3 | -16 | -5 | 10 | 270 | 2.46 |
| Rostral middle frontal | -21 | 47 | 31 | 26 | 702 | 2.59 |
| Frontal pole | -6 | 62 | -11 | 35 | 945 | 2.41 |
| Lateral occipital | -45 | -76 | -2 | 22 | 594 | 2.34 |
| Superior temporal | -48 | -1 | -5 | 17 | 459 | 2.3 |
| Fusiform | 27 | -49 | -20 | 13 | 351 | 2.16 |
| Lingual | 6 | -58 | 4 | 17 | 459 | 2.01 |

**Table 3:**
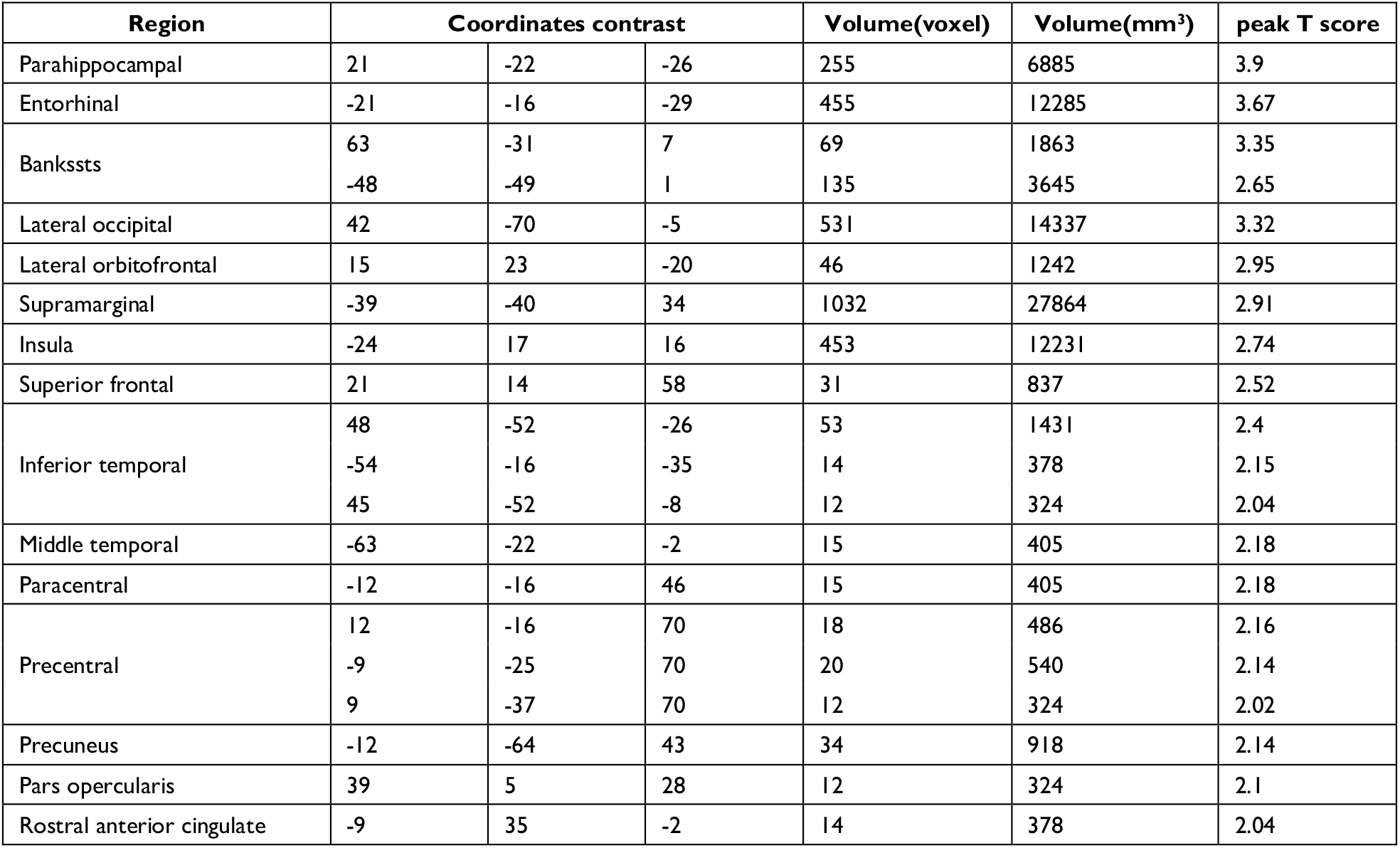
Regions showing negative BOLD-pain correlations.

| Region | Coordinates contrast |  |  | Volume(voxel) | Volume(mm<sup>3</sup>) | peak T score |
| --- | --- | --- | --- | --- | --- | --- |
| Parahippocampal | 21 | -22 | -26 | 255 | 6885 | 3.9 |
| Entorhinal | -21 | -16 | -29 | 455 | 12285 | 3.67 |
| Bankssts | 63 | -31 | 7 | 69 | 1863 | 3.35 |
|  | -48 | -49 | 1 | 135 | 3645 | 2.65 |
| Lateral occipital | 42 | -70 | -5 | 531 | 14337 | 3.32 |
| Lateral orbitofrontal | 15 | 23 | -20 | 46 | 1242 | 2.95 |
| Supramarginal | -39 | -40 | 34 | 1032 | 27864 | 2.91 |
| Insula | -24 | 17 | 16 | 453 | 12231 | 2.74 |
| Superior frontal | 21 | 14 | 58 | 31 | 837 | 2.52 |
| Inferior temporal | 48 | -52 | -26 | 53 | 1431 | 2.4 |
|  | -54 | -16 | -35 | 14 | 378 | 2.15 |
|  | 45 | -52 | -8 | 12 | 324 | 2.04 |
| Middle temporal | -63 | -22 | -2 | 15 | 405 | 2.18 |
| Paracentral | -12 | -16 | 46 | 15 | 405 | 2.18 |
| Precentral | 12 | -16 | 70 | 18 | 486 | 2.16 |
|  | -9 | -25 | 70 | 20 | 540 | 2.14 |
|  | 9 | -37 | 70 | 12 | 324 | 2.02 |
| Precuneus | -12 | -64 | 43 | 34 | 918 | 2.14 |
| Pars opercularis | 39 | 5 | 28 | 12 | 324 | 2.1 |
| Rostral anterior cingulate | -9 | 35 | -2 | 14 | 378 | 2.04 |

### Deep learning-based analysis with CNNs

A CNN was trained and applied to fMRI data to predict pain levels (see **Figure 3**). The prediction accuracy averaged across all eight-folds was 73% (**Figure 6 (A)**), which is significantly higher than the chance level of 50% (*P<2*10*^*-3*^). During the training phase, we recorded the value of the validation accuracy for each epoch, and as shown in **Figure 6 (B)**, the validation accuracy stabilized after 15 training epochs. According to the ROC curve in **Figure 6 (C)**, the area under the ROC curve (area under the curve (AUC)) for the low pain class was 70%, and the high pain class was 70%, demonstrating that the proposed CNN decoding approach could distinguish low pain and high pain levels via fMRI BOLD signals and detect more true positive and true negative samples than false positive and false negative samples.

**Figure 6.**
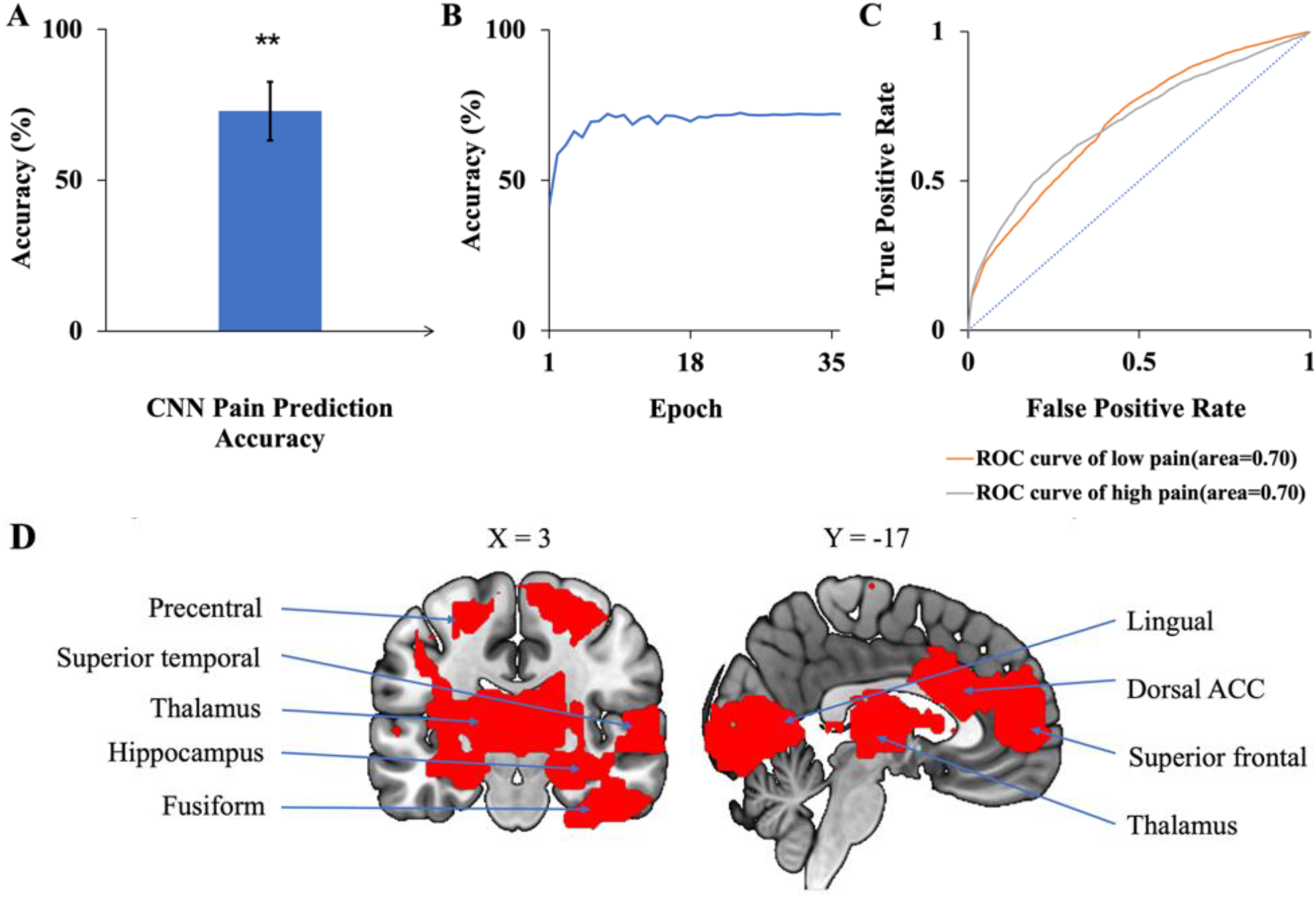
CNN analysis. **(A)** Accuracy of CNN prediction of pain levels is significantly above chance level of 50%. **(B)** Model prediction accuracy with learning. Prediction accuracy stabilized after 15 training epochs. **(C)** ROC curves where the AUC value for the low pain class was 0.70 while the AUC value for the high pain class was 0.70. **(D)** Brain regions contributing to CNN prediction performance, which included dACC, superior frontal, superior temporal, and lingual gyrus. \*\**P<2*10*^*-3*^

To identify the contribution of each brain region to the CNN classification of pain levels, we performed a sensitivity analysis of the trained CNN models by occluding the BOLD activities from each brain region and examined the decoding accuracy change without the contribution of the voxels in the region.^31^ A larger decrease in decoding accuracy is taken to indicate that the brain region being occluded plays a more important role in pain generation and perception. The important brain regions thus identified, as shown in **Figure 6 (D)**, included lingual, superior frontal, thalamus, and dorsal anterior cingulate cortex (dACC). A list of CNN-identified brain regions underlying TN pain is shown in **Table 8**.

### Deep learning-based analysis with GCNNs

A GCNN was trained to take fMRI functional connectivity patterns as input to predict pain levels. The prediction accuracy averaged across all eight-folds was 72% (**Figure 7 (A)**), which is significantly higher than the chance level of 50% (*P<2*10*^*-3*^). During the training phase, we recorded the value of the validation accuracy for each epoch, and as shown in **Figure 7 (B)**, after 20 epochs, the validation accuracy stabilized. According to the ROC curve in **Figure 7 (C)**, the area under the ROC curve (AUC) for the low pain class was 54%, and the high pain class was 66%, demonstrating that the GCNN approach worked well to decode low and high pain from function connection patterns.

**Figure 7.**
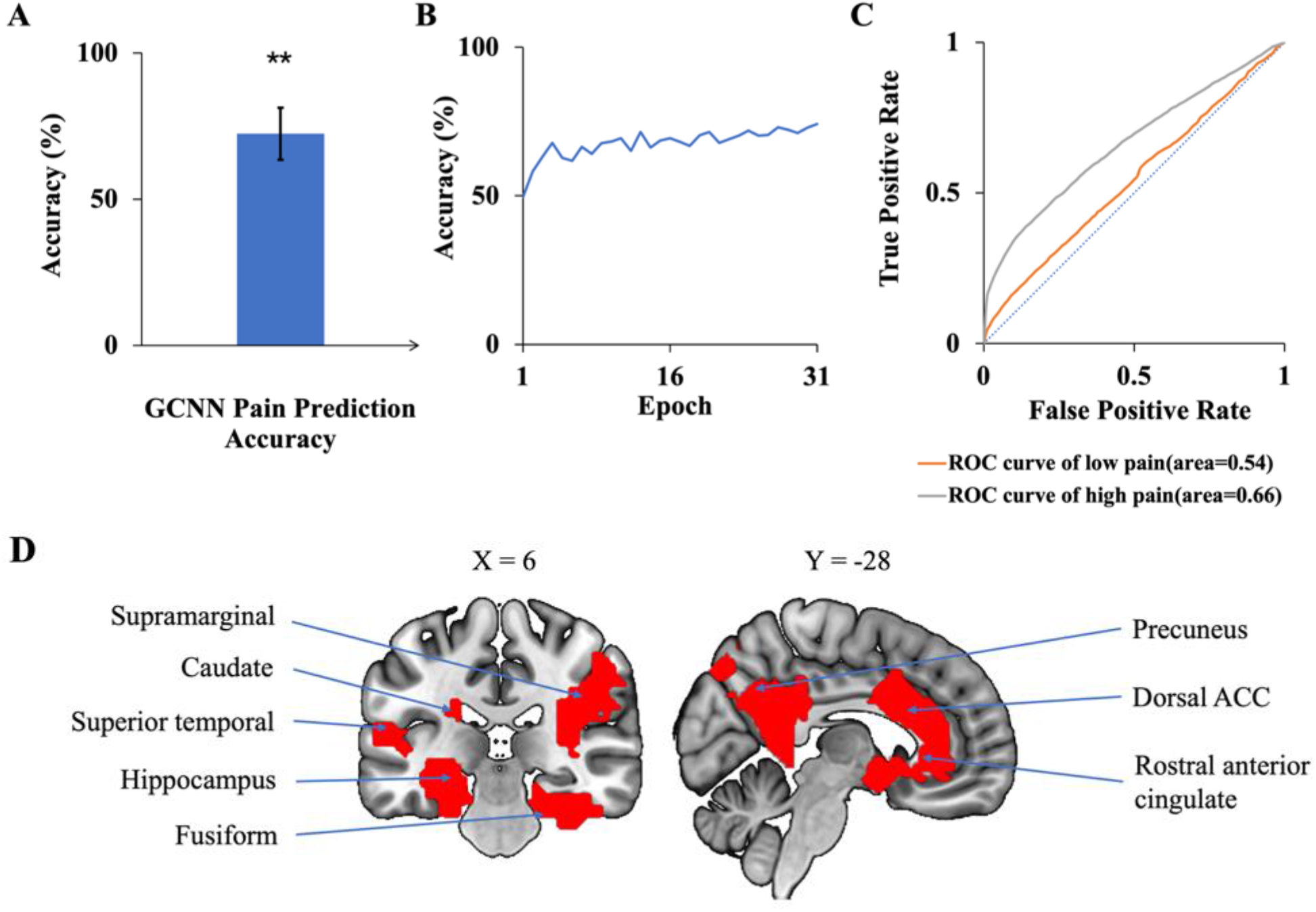
GCNN analysis. **(A)** Accuracy of GCNN prediction of pain levels is significantly above chance level of 50%. **(B)** Model prediction accuracy with learning. Prediction accuracy stabilized after 20 training epochs. **(C)** ROC curves where the AUC value for the low pain class was 0.54 while the AUC value for the high pain class was 0.66. **(D)** Brain regions contributing to GCNN prediction performance included dACC, fusiform, superior temporal, and precuneus. \*\**P<2*10*^*-3*^

We applied the same sensitivity analysis to the proposed GCNN model by occluding the BOLD activities and associated connections for each ROI. ROIs were ranked according to the prediction accuracy decrease from occlusion. Top-ranked brain regions, as shown in **Figure 7 (D)**, included dACC, fusiform, and superior temporal. A list of GCNN-identified brain regions underlying TN pain brain is shown in **Table 8**.

### Validation analyses

Both CNN and GCNN analyses are AI-inspired methods. While these methods have been successfully applied in a variety of fields,^32,33^ to date, they have not been extensively applied to analyze neuroimaging data. We performed two additional analyses to further test the validity of the two methods. First, it is reasonable to expect that the CNN and GCNN predicted pain levels during the pain tracking session and the patient’s self-reported pain levels be related. Assigning low pain the value of zero and high pain the value of one, the pain ratings predicted by CNN and GCNN based on fMRI averaged over the pain tracking session were plotted against patients’ reported pain ratings average over the pain tracking session in **Figure 8 (A)** and **8 (C)**, respectively, where a significantly positive correlation was seen for both CNN and GCNN (*R=0*.*62* and *0*.*60* with both *P<2*10*^*-3*^), suggesting that the CNN and GCNN predicted pain levels from fMRI brain scans indeed reflected the level of pain experienced by the patients. Second, as indicated in Methods, in our experimental paradigm, pain tracking and resting state were recorded back-to-back. It is reasonable to expect that the average pain levels during the two sessions are correlated. If the model-predicted pain levels indeed reflect the actual pain levels, a strong correlation between model-predicted pain during resting state and that during pain tracking should exist. This was found to be the case in **Figures 8 (B) and 8 (D)**.

**Figure 8.**
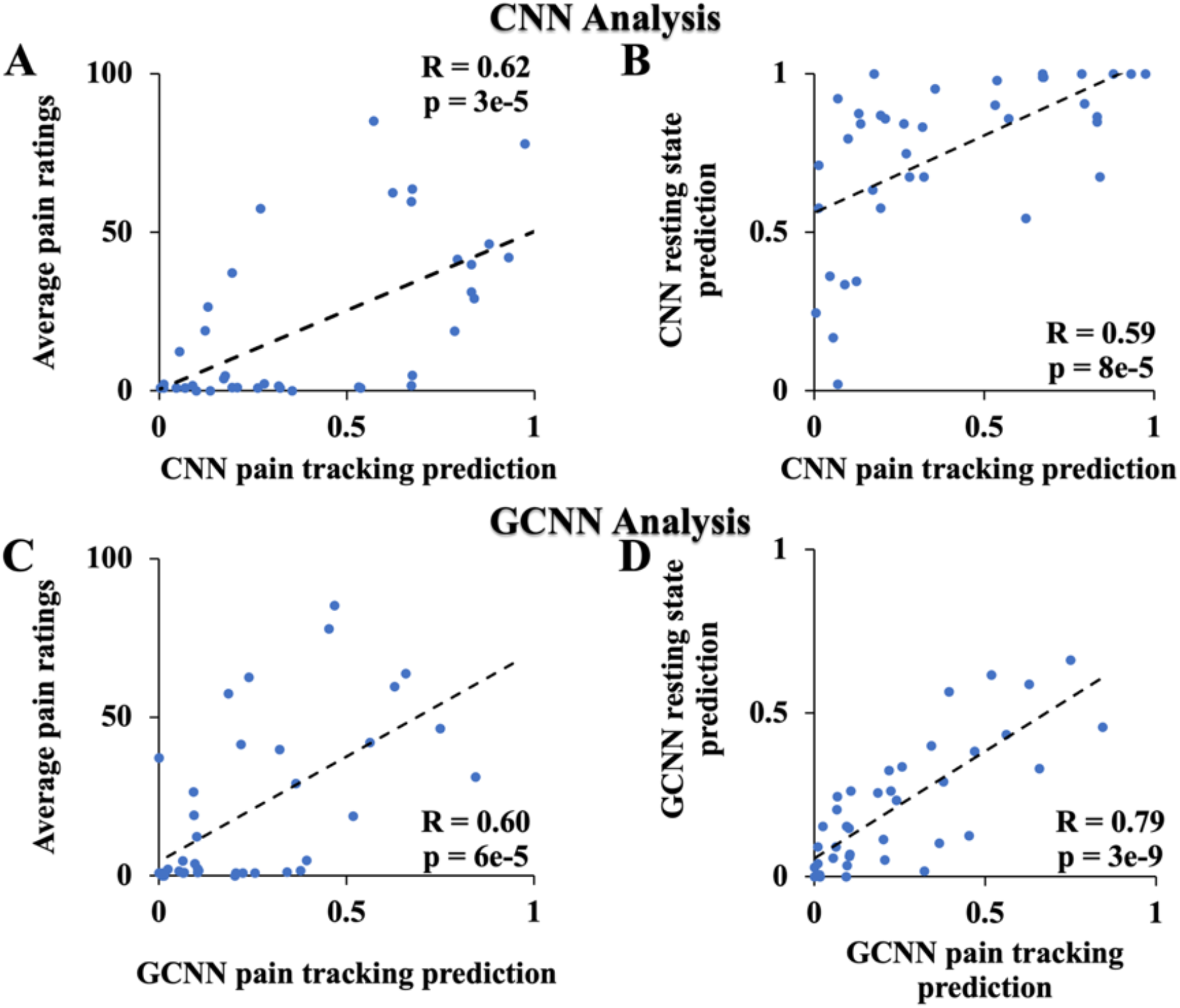
Additional CNN and GCNN analyses. **(A)** CNN predicted pain level vs average pain ratings. **(B)** CNN predicted pain level from resting state data vs. from pain tracking data. **(C)** GCNN predicted pain level vs average pain ratings. **(D)** GCNN predicted pain level from resting state data vs. from pain tracking data.

### Pain tracking vs motion tracking

As described in Methods, for the first 10 patients, 10-minutes of pain tracking was followed by 10 minutes of motion tracking. We tested, on the data from these 10 patients, whether the fMRI activities during pain tracking mainly reflected spontaneous pain fluctuations rather than movement executions. The BOLD time courses were extracted from each voxel in the pain tracking session (0-10 mins) and the motion tracking session (10-20 mins). We used the trained CNN models to compute the accuracy of pain rating predictions for the two sessions. The prediction accuracy for the pain tracking session was 68%, well above chance level of 50% (*P=0*.*04*), and that for the motion tracking session was 42%, which is not different from chance (*P=0*.*33*). This result was expected because the “pain level” during motion tracking was not the actual pain level experienced by the patient at the time of motion tracking.

### Subgroup analysis

Besides the male and female subgroups, among the 39 TN patients, there were 20 who had prior surgery for their conditions, but the pain returned at the time of the experiment. The remaining 19 patients never had surgery. A question of interest is whether these subgroups of patients have common neural substrate of pain generation and perception.

To test whether the subgroups shared common neural substrate of pain generation and perception, we trained the CNN and GCNN models on one subgroup and evaluated the performance of the model on the other subgroup. First, for models trained on female patients and tested on male patients, the decoding accuracy achieved by CNN and GCNN were 76.4% and 66.6% respectively, both significantly above chance level of 50% (both *P<2*10*^*-3*^). We did not train models on male patients to test them on female patients because the number of male patients is too small for model training purposes (*n=14* for male patients compared to *n=25* for female patients).

For models trained on non-surgery patients and tested on surgery patients, the decoding accuracy for CNN and GCNN were 55.7% and 53.3%, respectively, both significantly above chance level of 50% (both *P<2*10*^*-3*^), whereas for models trained on surgery patients and tested on non-surgery patients, he decoding accuracy achieved were 66% and 72% for CNN and GCNN models respectively, also both significantly above chance level of 50% (both *P<2*10*^*-3*^). See **Figure 9**. These results demonstrated that common neural substrate is shared among TN patients of different subgroups.

**Figure 9.**
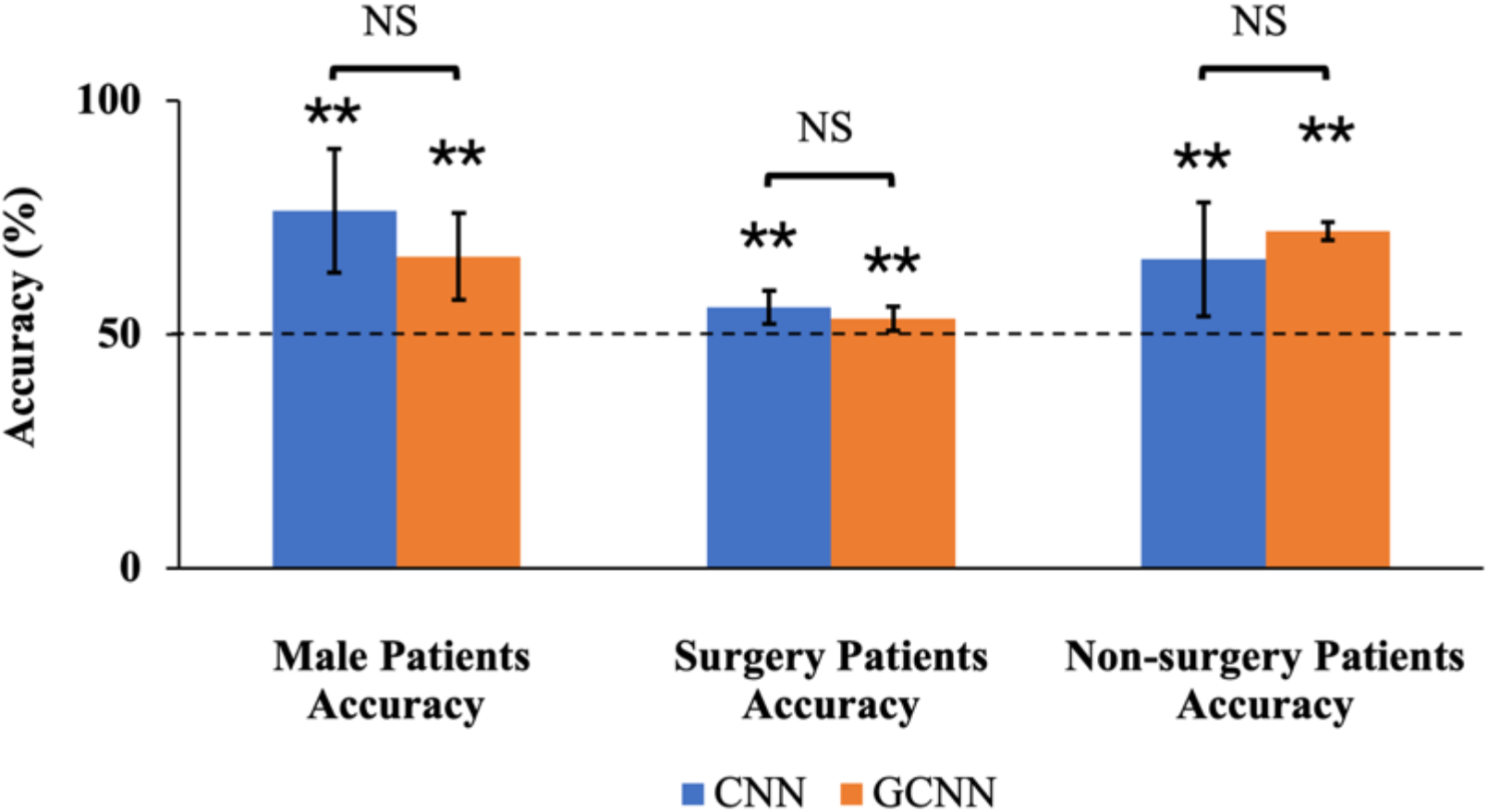
Sub-group Accuracies by CNN and GCNN. We trained CNN and GCNN models in different subgroups and tested the model performances in the remaining subgroup. By dividing the patients by gender, we found that both CNN and GCNN trained using female patients could achieve above-chance prediction accuracy on male patients. Dividing the patients into surgery/non-surgery subgroups, we found that CNN and GCNN models trained on one subgroup achieved above-chance decoding accuracy when tested on the other subgroup. NS: Not significant. \*\**P<2*10*^*-3*^

The clinical and psychological characteristics of the non-surgery vs. surgery and female vs. male subgroups were also evaluated. There were no significant differences between the surgery/non-surgery groups in age (*P=0*.*3848*), disease duration (*P=0*.*3593*), current pain intensity (*P=0*.*1027*), or usual pain intensity (*P=0*.*9055*). (**Figures 10-11**). When evaluating the different psychological outcomes from the different test inventories, there was a significant difference in the BDI-2 score, with surgical subjects having a significantly higher score (mean ± SD: 15±9 vs 9±8, *P=0*.*0487*). The other psychological scores from the PCS (*P=0*.*5237*), PASS (*P=0*.*3346*) (including subscales: fear (*P=0*.*8547*), cognitive (*P=0*.*4162*), escape/avoidance (*P=0*.*4071*), and physiological (*P=0*.*2511*)) were not significantly different between the surgery and non-surgery groups (**Tables 4-5**). When evaluating sex differences for these same clinical and psychological outcome measures, there was a significant sex difference in the age of the subjects, with males (mean ± SD: 65 ± 10 y.o.) being significantly older (*P=0*.*0025*) than female subjects (mean ± SD: 53±11). There were no significant differences between female and male subjects when comparing disease duration (*P=0*.*8997*), current pain intensity (*P=0*.*6229*), and usual pain intensity (*P=0*.*0759*) distributions (**Figures 12-13**). The psychological scores from the PCS (*P=0*.*5592*), BDI-2 (*P=0*.*5171*), and PASS (*P=0*.*5777*) (including subscales: fear (*P=0*.*7299*), cognitive (*P=0*.*9528*), escape/avoidance (*P=0*.*5393*), and physiological (*P=0*.*0776*)) tests were also not significantly different between female and male subjects (**Tables 6-7**).

**Figure 10.**
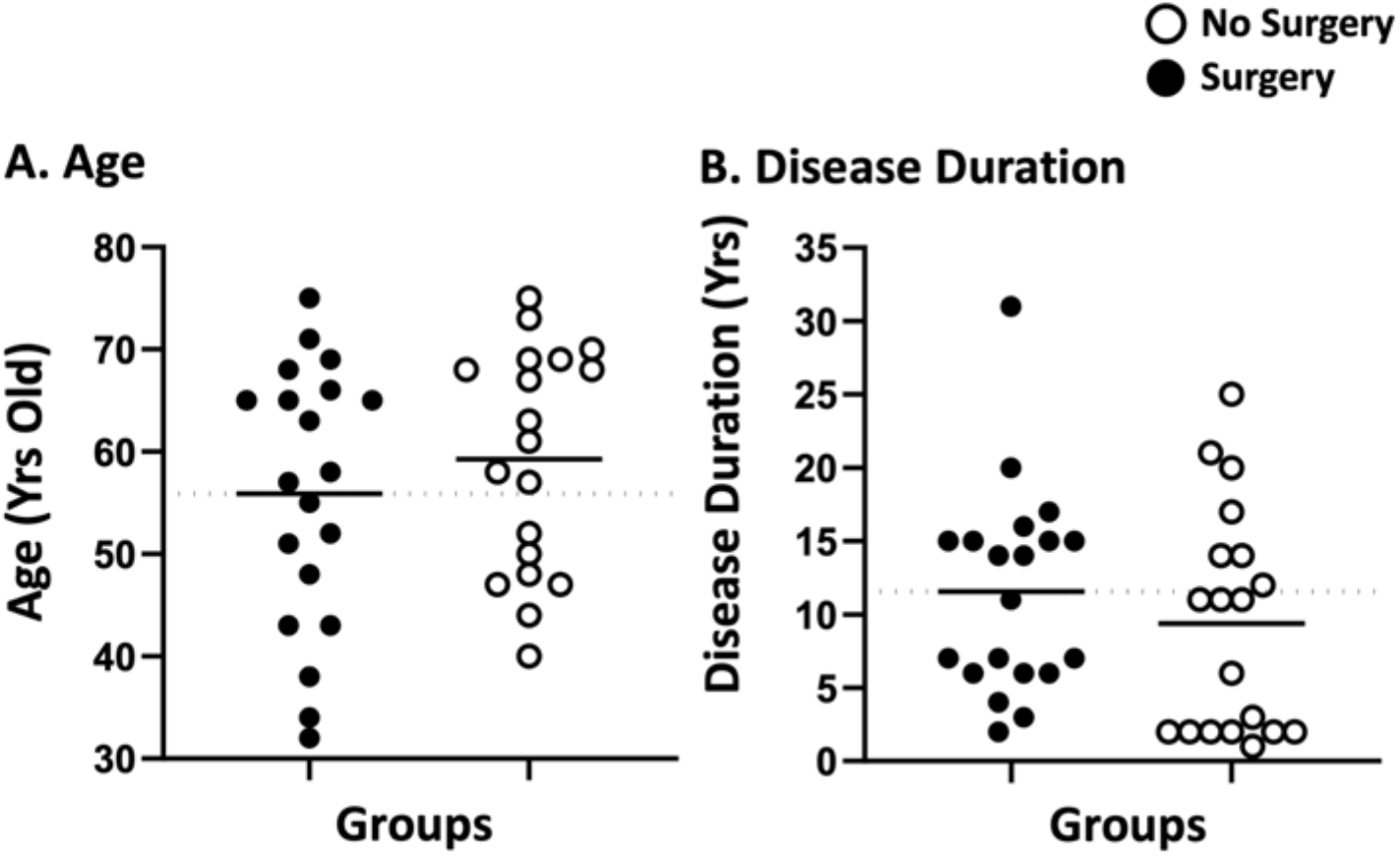
Surgery vs. non-surgery subgroup. **(A)** Age distribution and **(B)** Disease duration. Surgical and non-surgical groups had a similar age distribution and there was no significant effect from surgical status on disease duration.

**Figure 11.**
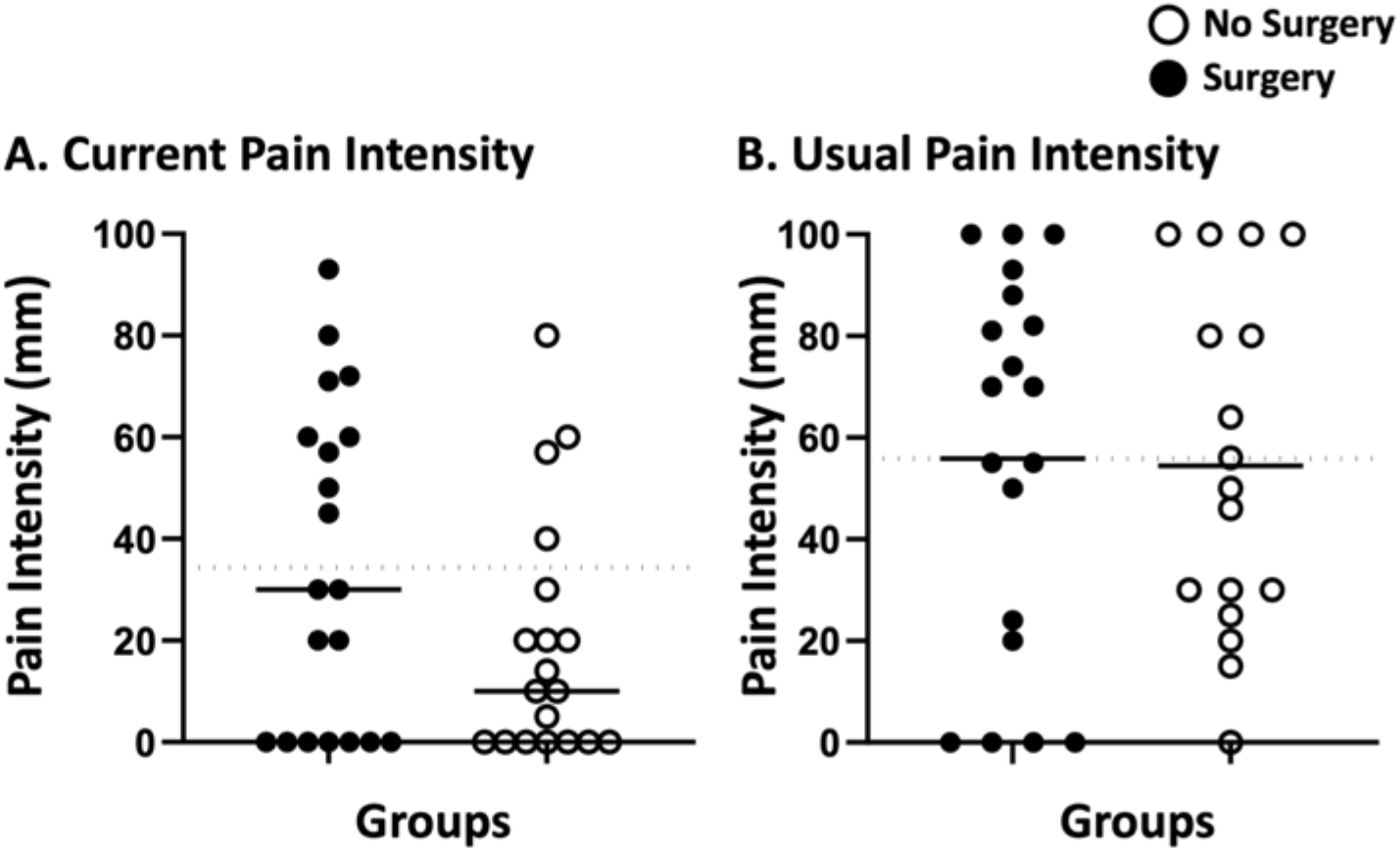
Surgery vs. non-surgery subgroup. **(A)** Current pain and **(B)** Usual pain intensity. Pain levels were similar for both surgical and non-surgical groups regarding the pain reported at time of scanning as well as reported usual pain intensity.

**Table 4.**
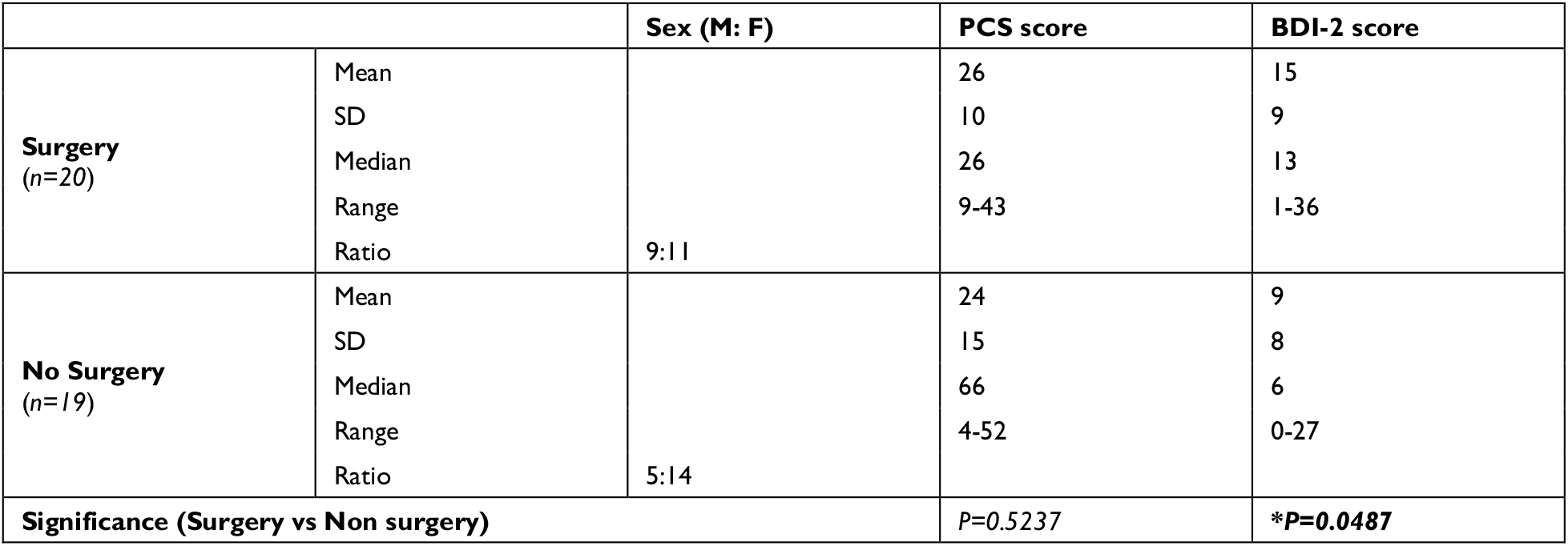
Surgery vs. non-surgery subgroup. Sex ratio, PCS, and BDI-2 information were shown. The BDI-2 score was significantly higher (*P=0*.*0487*, unpaired t-test) for the surgical subjects, indicating that this group likely suffered from higher depression as compared to subjects that did not have surgery. PCS scores were not significantly different between the two groups.

**Table 5.** Surgery vs. non-surgery subgroup. PASS total and subcategory scores were shown. The PASS total and subscale (Fear, Cognitive, Escape/Avoidance, and Physiological) scores were not significantly different between surgery and non-surgery subjects. This indicates that they exhibited similar levels of pain and anxiety feelings.

|  |  | PASS total score | PASS Factor 1: Fear | PASS Factor 2: Cognitive | PASS Factor 3: Escape/Avoidance | PASS Factor 4: Physiological |
| --- | --- | --- | --- | --- | --- | --- |
| **Surgery**<br>(n=20) | Mean | 90 | 21 | 28 | 26 | 16 |
|  | SD | 27 | 8 | 8 | 7 | 10 |
|  | Median | 87 | 20 | 26 | 27 | 14 |
|  | Range | 50-135 | 10-28 | 16-39 | 10-42 | 0-33 |
| **No Surgery**<br>(n=19) | Mean | 79 | 21 | 25 | 24 | 12 |
|  | SD | 41 | 10 | 11 | 12 | 12 |
|  | Median | 66 | 20 |  | 26 | 9 |
|  | Range | 12-170 | 6-40 |  | 1-43 | 0-50 |
| **Significance (Surgery vs Non surgery)** |  | *P*=0.3346 | *P*=0.8547 | *P*=0.4162 | *P*=0.4071 | *P*=0.2511 |

**Figure 12.**
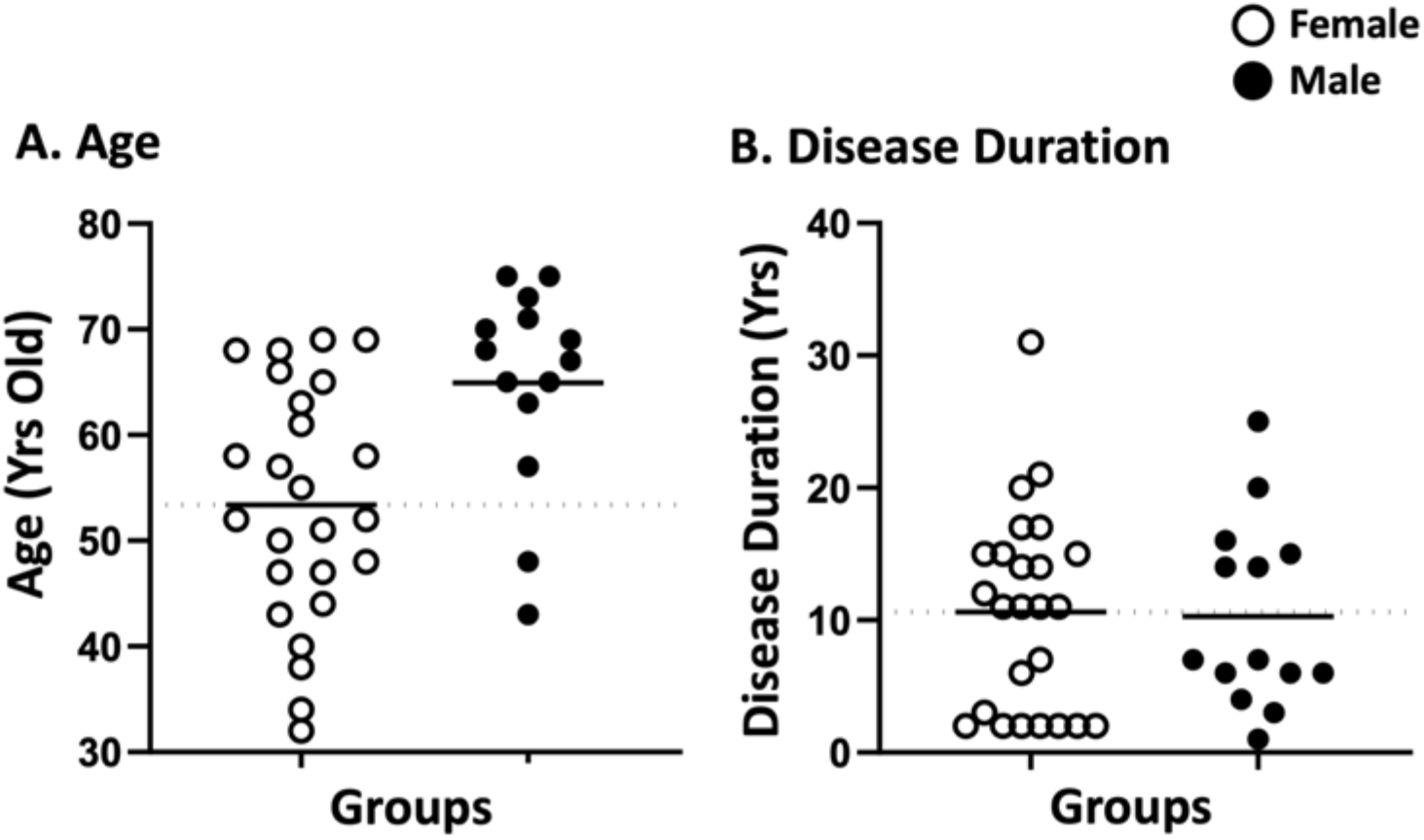
Female vs. male subgroup. **(A)** Age distribution and **(B)** Disease duration. Male subjects were significantly older (*P=0*.*0025*, unpaired t-test) as compared to female subjects; however, there was no difference in disease duration between the sexes.

**Figure 13.**
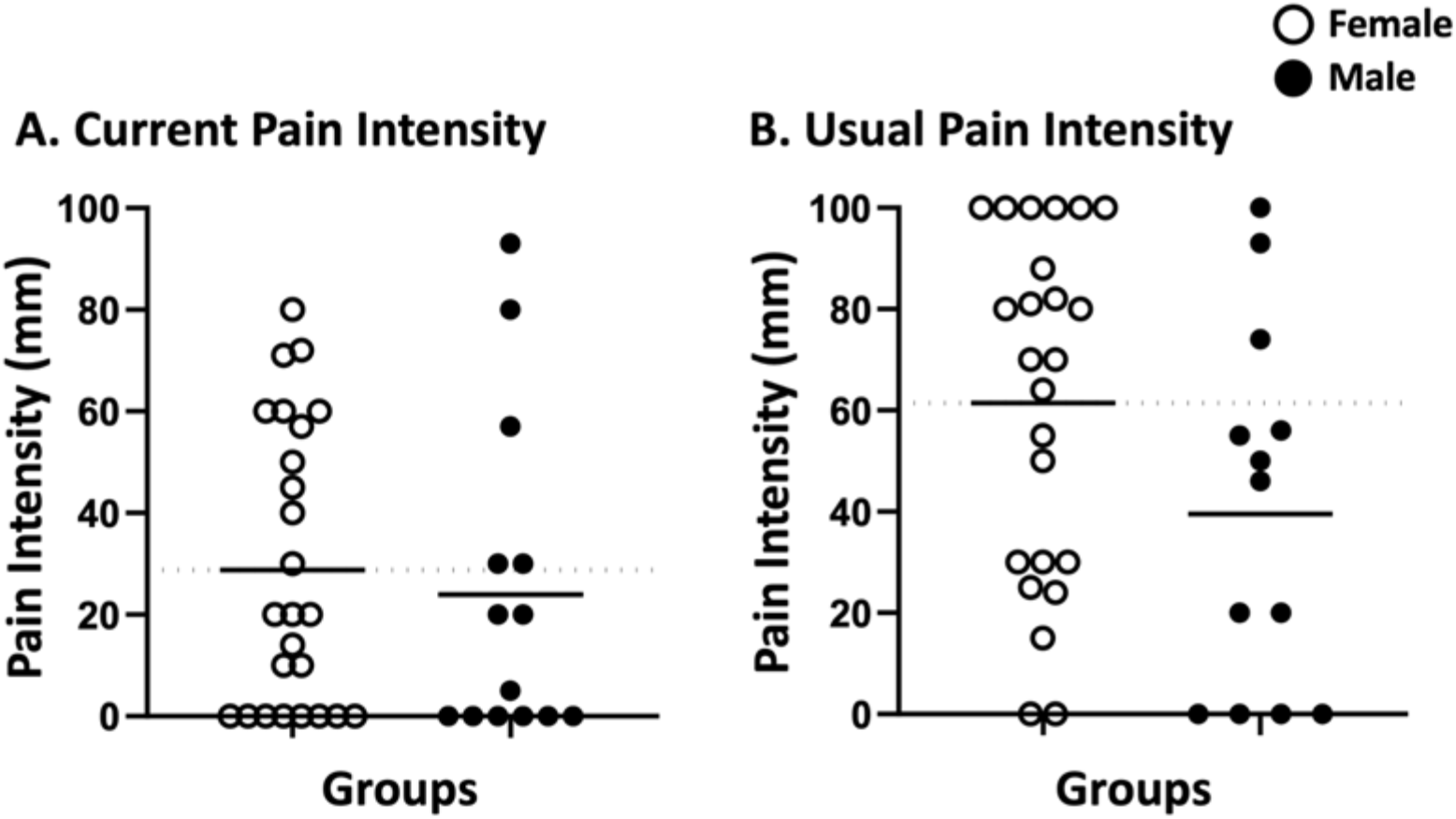
Female vs. male subgroup. **(A)** Current pain and **(B)** Usual pain intensity. There was no sex difference in either current or usual pain intensity.

**Table 6.** Female vs. male subgroup. PCS and BDI-2 information were shown. There was no sex difference in the PCS and BDI-2 test scores, indicating that female and male subjects exhibited similar catastrophizing and depression measures.

|  |  | PCS score | BDI-2 score |
| --- | --- | --- | --- |
| **Female**<br>(n=25) | Mean | 26 | 13 |
|  | SD | 13 | 10 |
|  | Median | 29 | 12 |
|  | Range | 4-52 | 0-36 |
| **Male**<br>(n=14) | Mean | 24 | 11 |
|  | SD | 11 | 7 |
|  | Median | 24 | 9 |
|  | Range | 7-36 | 3-23 |
| **Significance (Female vs Male)** |  | *P*=0.5592 | *P*=0.5171 |

**Table 7.**
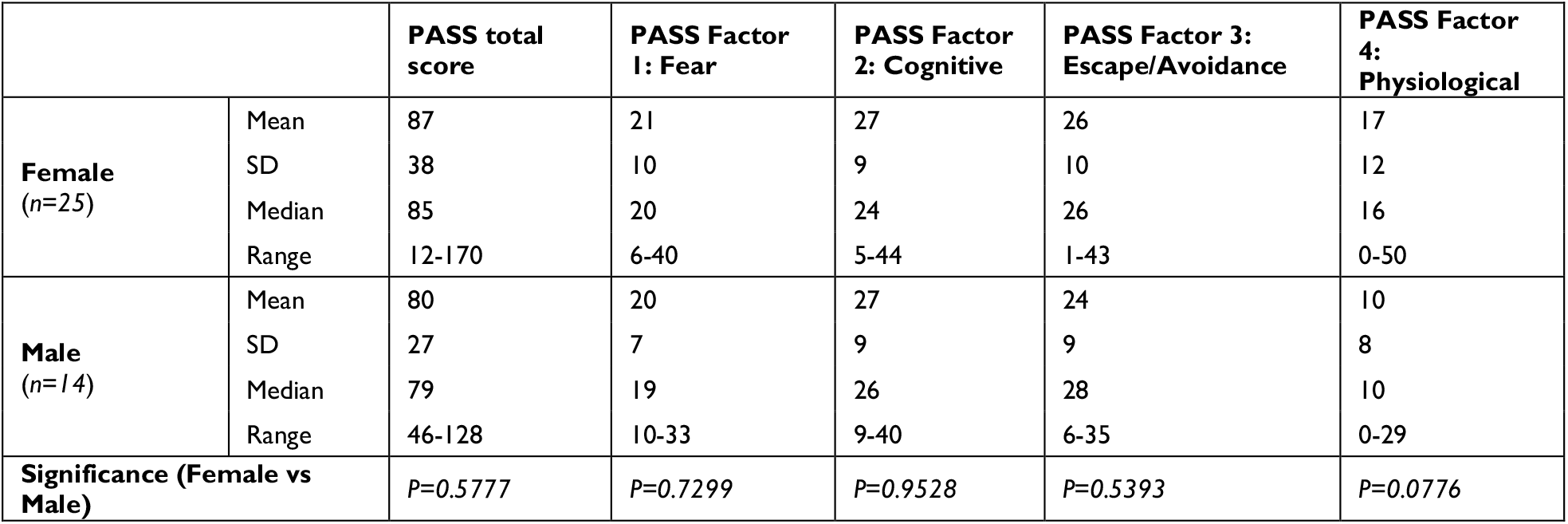
Female vs. male subgroup. PASS total and subcategory scores were shown. The PASS total and four subscale scores were not significantly different between males and females.

**Table 8:**
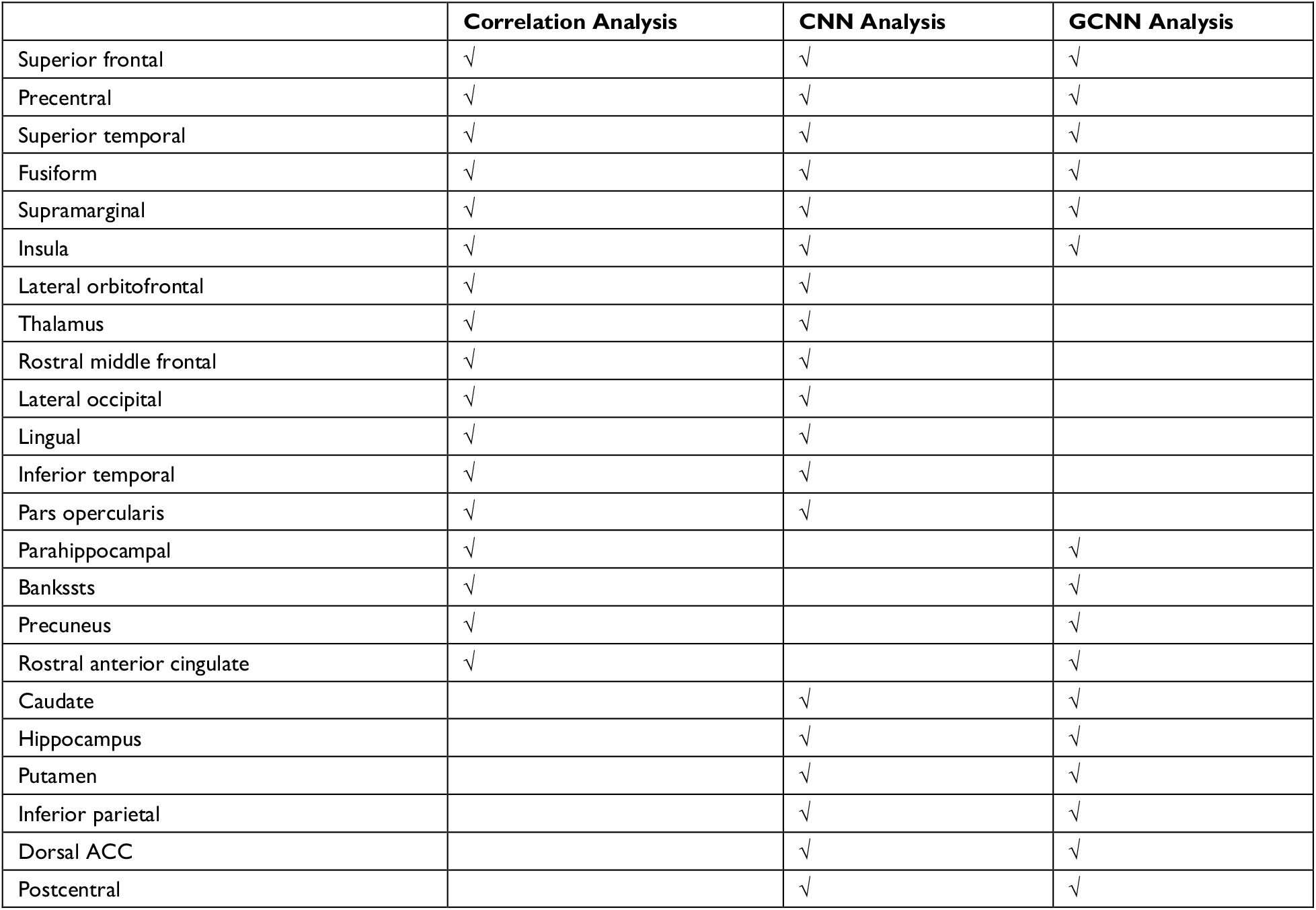
Signature centers of TN pain identified by correlation and AI-inspired analyses.

### Signature centers of TN pain

Three analyses were applied in this study to identify signature centers of TN pain: correlation analysis, CNN analysis, and GCNN analysis. We listed the top 23 brain regions that appeared in at least two of the analyses in **Table 8**. Six regions were identified by all three methods, including superior temporal, insula, fusiform, precentral gyrus, superior frontal gyrus, and supramarginal gyrus. The 17 remaining regions included dACC, thalamus, lateral occipital gyrus, inferior temporal, postcentral gyrus, lingual gyrus, and inferior parietal gyrus. All these regions have been implicated in chronic pain especially TN pain in past studies (see Discussion).

## Discussion

In this study we sought to identify the neural substrate of TN pain by recording and analyzing fMRI data from TN patients while they rested or tracked their spontaneous fluctuations in pain levels. By applying both conventional and AI-inspired approaches, we obtained converging evidence implicating a common set of brain regions, including the insula and the precentral gyrus, as playing an important role in the generation and perception of TN pain. Additional complementary insights into TN pain-related brain regions are offered by each of the three approaches.

### Signature pain centers identified by three analyses

**Table 8** shows that there are six brain regions that appeared in all three analyses, including superior temporal, insula, fusiform gyrus, precentral gyrus, superior frontal gyrus, and supramarginal gyrus. All these regions have been implicated in TN pain in prior studies.^34^ Yuan et al. and Xiang et al.^7,35^ reported increased ReHo in the superior frontal gyrus in TN patients compared to healthy controls; cortical thickness was shown to decrease with increase in pain duration.^36^ The superior temporal gyrus (STG), functionally known for its role in memory and language processing,^37^ showed gray matter volume abnormalities in TN, and the degree of grey matter volume alterations in the left STG may reflect pain severity. The abnormal structural alterations in the temporal lobe may be due to the generation and/or maintenance of emotions and perception of pathological pain.^38^ The important role of precentral and postcentral gyrus in pain processing is well established. The primary motor cortex, located in the precentral gyrus, may be involved in inhibiting motion so as not to aggravate pain conditions,^39^ whereas the postcentral gyrus, where the primary somatosensory cortex is located, is the cortical gateway through which nociceptive input is processed and transmitted to other brain structures.^40-43^ In a resting-state fMRI study, Wang et al.^9^ found that ReHo in the precentral gyrus and the postcentral gyrus is positively correlated with patients’ daily experienced pain severity, suggesting a link between local synchronization of intrinsic brain activity and pain modulation. In addition, in a recent meta-analysis, the somatosensory cortex is found to be structurally changed in TN compared with healthy controls.^44^ The fusiform gyrus, a visual area known for its involvement in face-related processing, is thought to be important in mediating mental imagery processes related to pain perception.^45,46^ In TN, the fusiform is shown to have decreased gray matter volume compared to controls.^47,48^ In addition, the left fusiform gyrus is found to be involved in pain anticipation and perception, leading to a negative correlation with pain ratings.^45^ The insular cortex has also been implicated in mediating pain intensity as well as negative emotions.^49^ A recent MRI study found altered insular morphology and functional connectivity (FC) and abnormal diffusion parameter in the white matter adjacent to the insular cortex in TN.^50^ Furthermore, significant local gyrification index (LGI) reductions in the left insular cortex were found in patients with TN compared with control groups.^50^ The authors argued that pain perception results from nociceptive representation being transformed into subjective magnitude assessment within the insula.^51^ Meanwhile, Tian et al.^52^ found that TN patients exhibited significantly increased long-range functional connectivity density in the right supramarginal gyrus. As can be seen, these past studies rely either on either structural information or resting state data. The functional roles of these regions in TN pain generation and perception remain to be better understood. Our results, by utilizing a functional paradigm in which TN patients tracked pain levels, shed new light on this issue. In particular, we found that the neural activities in these six regions not only closed tracked the pain level fluctuations (correlation analysis and CNN analysis), they also mediated network-level communications among different brain regions (GCNN analysis).

### Signature pain centers identified by two of the three analyses

Among the common set of regions identified by both AI-inspired analyses but not the correlation analysis, dACC is known for its role in pain processing^44^; it is consistently activated in human imaging studies of pain. As a central hub in the pain matrix, the dACC is highly connected to other brain areas involved in cognition, emotion, and negative affect, all of which are associated with chronic pain.^53-55^ It has been further suggested that the cingulate cortex is also important for the transition from acute to chronic pain.^56^ Previous TN works, including the Yuan et al.’s^7^ resting-state fMRI data and Moisset et al.’s^12^ task fMRI data, have found abnormal activities in ACC. Structurally, Mo et al.^57^ found that TN patients exhibited reductions in cortical indices in the ACC, the midcingulate cortex (MCC), and the posterior cingulate cortex (PCC) relative to healthy controls group, indicating that the ACC may play a role in pain adaptation, habituation, distraction, and the engagement of the endogenous pain control system.^58^ In addition to dACC, the thalamus, the postcentral, the lingual, and the inferior temporal gyrus are found to be activated in two of the three analyses. These regions have also been implicated in previous research on TN pain. Previous research has firmly established the importance of the thalamus in pain processing; it receives nociceptive sensory information from the periphery, integrates this information with arousal and attention, and sends outputs to broad regions of the cerebral cortex for further processing.^59-61^ In TN, the grey matter increases in the thalamus for TN patients relative to controls, suggesting a link between thalamic structural change and activity-dependent plasticity in S1 via thalamocortical projections.^49,62^ In our data, the GCNN analysis was not able to show that the thalamus is an important region underlying TN pain, whereas the correlation and CNN methods found that the thalamus is important for predicting the spontaneous fluctuations of TN pain, consistent with the established role of the thalamus in pain processing including TN pain.^63^ Regarding the inferior temporal gyrus, past work found that the gray matter volume of the left inferior temporal gyrus was negatively correlated with current pain intensity and disease duration in TN patients.^48,64^ Decreases in the amplitude of low-frequency fluctuation (ALFF) in the right inferior temporal region were found in TN patients.^65^ Zhu et al.^11^ also found that compared with the healthy control (HC) group, patients with TN showed the degree centrality value (calculated by counting the number of significant suprathreshold correlations (the degree of the binarized adjacency matrix) for each individual) changed in the right lingual gyrus, right postcentral gyrus, left paracentral lobule, left inferior cerebellum, and right inferior cerebellum.

### Clinical considerations

The clinical TN population that was recruited for this study was consistent with the known disease demographics and characteristics: female > male and age > 50 years old.^66^ The majority of subjects (*n=16*) reported moderate to severe usual pain levels, seven of which reported maximal VAS pain levels of 100 (**Figure 11 (B)**). At the time of the scanning procedure, the current pain intensity was distributed roughly equally across zero to maximum pain range, indicating the spontaneous nature of this pain disorder. There was a significant age difference between the males and females; however, with the males being on average older; the disease duration was similar between males and females. This likely indicates that males were diagnosed later in life as compared to females. The only psychological factor that was elevated was the BDI-2 score for surgical subjects, as compared to non-surgical subjects. While significant (*P=0*.*0487*), the average male BDI-2 score is considered within the “mild-to-moderate depression” range (mean ± SD: 15±9), while the female scores were within the no or minimal depression range (mean ± SD: 9±8). On average, the PCS ratings for the subjects are within the mild range (21-40), and the PASS scores were not significantly different when comparing surgical status or sex differences. In the context of these clinical findings, our results that both CNN and GCNN models constructed based on patients from one subgroup (surgical vs. non-surgical, male vs. female) can decode the patients from the other subgroup can be seen as reflecting shared neural substrate rather than driven by differences in clinical conditions.

### Methodological considerations

The correlation analysis has been used in numerous pain studies and remains the principal method of percept-related fMRI for identifying the neural substrate of naturally occurring pain.^15^ It is simple and intuitive.^49^ The weaknesses are that it only detects linear relationships, and the statistical effects are not strong. In our data, if we applied any kind of multiple comparison correction, we would find no activations. Deep learning is an emerging area of machine learning, and it is in the early stages of being applied to analyze neuroimaging data. In this work, in addition to showing that CNN and GCNN are able to predict pain fluctuations from fMRI data, we did several additional analyses to validate the approach. First, we showed that the predicted pain levels by CNN and GCNN are correlated with the pain ratings indicated by the patients, as would be expected. Second, the CNN and GCNN predicted pain levels from the resting-state data and from pain tracking data are highly correlated, again as would be expected, demonstrating that the CNN and GCNN model predictions are robust. Combining these AI-inspired methods with conventional method to seek converging evidence may become a promising way in future studies of pain neuroimaging.

### Limitations

This study has a number of limitations. First, for the correlation analysis, as mentioned above, if a whole-brain multiple comparison approach such as false discovery rate (FDR) was applied, no statistically significant regions will appear in the activation map, despite having a reasonable sample size of *n=39*. The situation is similar in prior pain studies utilizing the percept-related fMRI analysis.^67^ Nevertheless, the thalamus, an established region in pain processing including TN pain, only appeared in the CNN and correlation analysis map.^68,69^ This demonstrates that, despite the statistical weakness associated with the correlation analysis, it can still provide important information which not all AI-inspired methods can provide. Second, because of the need to increase the pain tracking session length without increasing the overall scanning time, we were only able to record motion tracking from the first 10 patients. Nevertheless, by applying the AI-inspired models to the data from pain tracking and from motion tracking in these 10 patients, we were able to establish that the fMRI data recorded during pain tracking are mainly driven by spontaneous pain fluctuations, not by the movement associated with pain tracking. Last, when assigning discrete class labels to the data, the threshold of 15 was chosen to achieve the class balance between low and high pain. This high vs. low pain level demarcation was a trade-off between clinical considerations and technical requirements and is not a clinically applicable pain level designation.

## Conclusion

We applied advanced statistical methods to patterns of brain activation related to the paroxysms of TN pain and generated a set of “signature centers” of pain generation and perception within the brain. Our approach, combining both conventional and AI-inspired methods to yield converging findings, is highly novel in the context of TN research, and our result provides a sorely needed basis for understanding the central mechanisms of TN. We hope that the insights revealed in this study can lead to a better understanding of TN and curing of this devastating condition.

## Abbreviations

AI: artificial intelligence
ALFF: amplitude of low-frequency fluctuation
ASA: American Society of Anesthesiologists
AUC: area under the curve
BDI-2: Beck Depression Inventory-II
BOLD: blood-oxygen-level-dependent
CBP: chronic back pain
CNN: convolutional neural networks
dACC: dorsal anterior cingulate cortex
DNN: deep neural network
EPI: echo-planar imaging
FC: functional connectivity
FDR: false discovery rate
fMRI: functional magnetic resonance imaging
FOV: field of view
FPRF: Facial Pain Research Foundation
GCNN: graph convolutional neural network
HC: healthy control
HRF: hemodynamic response function
IBS: irritable bowel syndrome
ICHD-3: the 3rd edition of the International Classification of Headache Disorders
IHS: International Headache Society
IRB: Institutional Review Board
LGI: local gyrification index
LR: logistic regression
MCC: midcingulate cortex
MVD: microvascular decompression
OHSU: Oregon Health Science University
PASS: Pain Anxiety Screening Scale
PCC: posterior cingulate cortex
PCS: Pain Catastrophizing Scale
ReHo: regional homogeneity
ROC: receiver operating characteristics
ROI: region of interest
rs-fMRI: resting-state fMRI
SD: standard deviation
SGD: stochastic gradient descent
SPM: statistical parametric mapping
SVM: support vector machine
TE: time of echo
TN: trigeminal neuralgia
TR: repetition time
VAS: visual analog scale

## Competing interests

The authors declare no conflicts of interest.

## Funding

This study was supported by the Facial Pain Research Foundation (FPRF).

## Notes

### Competing Interest Statement

The authors have declared no competing interest.

